# Mitochondrial translation is a targetable dependency of chemo-refractory triple negative breast cancer

**DOI:** 10.64898/2026.03.23.712647

**Authors:** Mariah J. Berner, Steven W. Wall, Mokryun L. Baek, Audra Lane, Allison S. Greer, Karen Wang, Lacey E. Dobrolecki, Ivy Strope, Qian Zhu, Bing Zhang, Jonathan T. Lei, Michael T. Lewis, Gloria V. Echeverria

**Affiliations:** Lester and Sue Smith Breast Cancer, Baylor College of Medicine, Houston, TX, USA; Dan L Duncan Comprehensive Cancer Center, Baylor College of Medicine, Houston, TX, USA; Department of Medicine, Baylor College of Medicine, Houston, TX, USA; Department of Molecular and Human Genetics, Baylor College of Medicine, Houston, TX, USA; Department of Molecular and Cellular Biology, Baylor College of Medicine, Houston, TX, USA; Department of Radiology, Baylor College of Medicine, Houston, TX, USA; Department of Radiation Oncology, Baylor College of Medicine, Houston, TX, USA

## Abstract

Triple negative breast cancer (TNBC) patients harboring residual cancer burden following completion of conventional neoadjuvant chemo-immunotherapy regimens have poor relapse-free and overall survival rates and limited therapeutic options. We and others have demonstrated that mitochondrial function is required for the survival of chemo-refractory TNBC. Here, we define the mitochondrial translation machinery as a critical and targetable dependency underlying chemo-refractory TNBC. Analyses of human and orthotopic patient-derived xenograft (PDX) mass spectrometry proteomics datasets revealed that mitochondrial protein translation-related signatures were among the top associated with chemoresistance. These signatures included core mitoribosome components and the mitoribosome-associated factor Oxidase (Cytochrome C) Assembly 1-Like (OXA1L), which was consistently enriched in chemoresistant versus chemosensitive TNBC across datasets. OXA1L, a key mediator of mitochondrial translation and electron transport chain (ETC) assembly, has not been functionally characterized in cancer. We therefore tested whether OXA1L-dependent mitochondrial translation sustains mitochondrial function and chemoresistance in TNBC. Knockdown (KD) of OXA1L in human TNBC cells reduced ETC protein levels, mitochondrial ‘respirasome’ supercomplex levels, ATP production, and oxidative phosphorylation (oxphos), establishing a requirement for OXA1L in maintaining mitochondrial bioenergetics in TNBC. OXA1L was required for the characteristic oxphos elevation induced by carboplatin (CRB), and KD significantly enhanced CRB sensitivity, demonstrating that mitochondrial translation supports adaptive metabolic responses to chemotherapy. To explore the translational potential of targeting the mitoribosome in TNBC, we leveraged the bacterial ancestry of mitochondria to repurpose the FDA-approved antibiotic tigecycline (TIG) as a mitochondrial translation inhibitor. Direct measurement of mitochondrial nascent peptide levels revealed that, while CRB elevated mitochondrial translation, TIG potently suppressed mitochondrial translation as monotherapy and in combination with CRB or docetaxel (DTX). TIG abolished CRB-induced oxphos, decreased oxphos in combination with DTX, and significantly improved chemotherapy sensitivity in human TNBC cell lines, PDX-derived spheroids, and *in vivo*. TIG sensitivity associated with mitochondrial translation-related proteomic signatures, concordant with PDX and patient-derived signatures associated with chemoresistance. Together, these data identify OXA1L-dependent mitochondrial translation as a targetable dependency that sustains mitochondrial function and chemoresistance in TNBC, demonstrate that its inhibition enhances chemotherapeutic response, and nominate a mitochondrial translation-related protein signature as a candidate predictive biomarker of TIG sensitivity and chemoresistance. These findings support mitochondrial translation inhibition as a potential therapeutic strategy in chemo-refractory disease.

**DISCLOSURES:** GVE is co-founder, Chief Scientific Officer, and an equity stakeholder of Nemea Therapeutics, Inc. G.V.E. formerly received sponsored research funding from Chimerix Inc. G.V.E. receives experimental compounds from the Lead Discovery Center of Germany and from Jazz Pharmaceuticals. MLB is a co-inventor at Nemea Therapeutics. MTL is a founder and limited partner in StemMed Ltd. and a manager in StemMed Holdings, its general partner. MTL is a founder and equity stakeholder in Tvardi Therapeutics Inc. The BCM PDX models, are exclusively licensed to StemMed Ltd., resulting in royalty income to MTL when used for commercial purposes. LED is a compensated employee of StemMed Ltd; however, the PDX models resulting in royalty income to LED were not included in this study. All other authors have nothing to disclose.

## INTRODUCTION

Triple negative breast cancer (TNBC) is an aggressive subtype lacking molecularly targeted therapy options for the 75-85% of patients lacking deleterious BRCA1/2 aberrations^1,2^. TNBC is defined by the molecular features it lacks, namely expression of the estrogen receptor, progesterone receptor, and overexpression or amplification of human epidermal growth factor receptor 2. The mainstay treatment for TNBC is a three to six-month course of neoadjuvant chemotherapy (NACT) consisting of several chemotherapeutic agents administered simultaneously and/or sequentially prior to surgery^1,3^. There is no one universally accepted NACT regimen^4^. Standard of care chemotherapies include agents that induce various forms of DNA damage (*e.g.* carboplatin (CRB), Adriamycin (aka doxorubicin), and cyclophosphamide) and microtubule-perturbing taxanes (*e.g.* docetaxel (DTX) and paclitaxel). NACT regimens were recently updated to include the immunotherapy agent Programmed Cell Death 1 (PD-1) inhibitor, with a modest yet statistically significant improvement in response rate^5^. Despite these many, lengthy treatments, residual tumors persist in approximately 45% of patients^6,7^. Presence of residual tumor burden following NACT is a strong and significant correlate of poor relapse-free and overall survival^6^. The five-year survival rate of metastatic TNBC is an unacceptable 12%^8^. Thus, identifying the molecular dependencies of tumor cells that evade NACT is of utmost importance to improve outcomes for women living with TNBC.

Mitochondrial metabolism has been reported to support chemotherapy resistance in hematologic malignancies and solid tumors of breast, colon, prostate, pancreas, and other sites^9–15^. We and others have established the importance of mitochondrial oxidative phosphorylation (oxphos) in TNBC^9,10,16–19^. Multi-omic analyses of cohorts of TNBC patients^19,20^ and orthotopic patient-derived xenograft (PDX) models^18^ have revealed that not only is oxphos the most upregulated metabolic pathway in TNBC relative to normal mammary tissue^20^, but also that oxphos is among the pathways most significantly and reproducibly associated with chemoresistance. We demonstrated that inhibition of mitochondrial electron transport chain (ETC) complex I with IACS-010759^9,21^ or of mitochondrial fusion protein optic atrophy 1 with MYLS22^10,22^ were both efficacious preclinical approaches to delay TNBC relapse, although the mechanistic underpinnings of mitochondrial dependence in TNBC remain poorly understood.

The mitochondrial genome encodes 37 genes, including 2 rRNAs, 22 tRNAs, and 13 mRNAs. Mitochondria have their own replication, transcription, and translation machinery. The mitoribosome translates the 13 mtDNA-encoded proteins, all components of ETC complexes I, III, IV, and V^23,24^. The remaining 79 ETC proteins are encoded in the nucleus (nDNA). Oxidase (Cytochrome C) Assembly 1-Like (OXA1L) is a transmembrane mitoribosome accessory protein insertase that has been shown in yeast and mammalian cells to interact with the mitoribosome via its C-terminal tail, promoting translation termination and insertion of nascent proteins into the inner mitochondrial membrane (IMM), where the ETC resides^24–26^. Furthermore, OXA1L is responsible for the insertion of 81% of the nuclear-encoded ETC subunits, initially transported into the matrix, into the IMM for the ETC to properly form^27^. Previous studies have demonstrated the importance of OXA1L for the formation and function of Complexes I, IV, and V^27–29^ in yeast and several human cell types, including skeletal muscle, dermal fibroblasts, and induced pluripotent stem cells. Its potential role in the function of Complexes II and III has not been established. OXA1L variants and polymorphisms have been linked with several mitochondrial disorders^29^, including childhood-onset mitochondrial encephalopathy^27^, asthma^30,31^, and mitochondrial myopathy^32^. Disruption of OXA1L function contributes to mitochondrial dysfunction in Parkinson’s Disease^33^ and in mitochondrial myopathy^32^. However, the functional relevance, regulation, and activity of OXA1L in cancer has not been reported.

Herein, we identified mitochondrial translation as a promising new, targetable metabolic vulnerability of chemoresistant TNBC, supported by correlative analyses of patient and PDX cohorts and functional studies in laboratory models. Analyses of multi-omic datasets^19,34,35^ revealed a pervasive, positive association of levels of mitochondrial ribosome proteins, as well as the accessory protein OXA1L, with TNBC chemoresistance. We provide novel evidence that OXA1L supports protein levels spanning all five ETC complexes. Further, we find mitochondrial translation and OXA1L are functional metabolic dependencies of chemoresistant TNBC. We went on to demonstrate that repurposing conventional antibiotics to inhibit mitoribosome function may be a promising approach to combat mitochondrial adaptations and survival of chemo-refractory TNBC cells.

## METHODS

### Analysis of publicly available datasets for molecular profiles of patient TNBC and PDX TNBC tumors

Quantitative chemotherapy responses measured as log_2_ fold change in tumor volume, and qualitative mRECIST categorization after four weekly cycles of single-agent carboplatin or docetaxel, along with log_10_ transformed mass spectrometry-based proteomic data from 50 TNBC PDXs, was downloaded from supplemental tables in the Lei et al, 2025 manuscript^18^. Associations between proteomics data and quantitative responses were assessed by Spearman correlation and Student’s t-test was used for proteomics data and qualitative responses. Log_2_ transformed Tandem Mass Tag (TMT) mass spectrometry-based proteomics data for the CPTAC-TNBC study was downloaded from supplemental tables in the Anurag et al, 2022 manuscript^19^. Baseline proteomic data was available for 48 patients treated with neoadjuvant carboplatin plus docetaxel (17 pCR, 31 non-pCR), including 14 paired samples collected after 48-72 hours of chemotherapy. For baseline analysis, Wilcoxon -rank-sum test was used to compare proteins of non-pCR samples to pCR samples. Gene Set Enrichment analysis with Reactome and MitoCarta geneset collections was performed using WebGestalt^36^ with signed -log_10_ p-values as input (positive sign if higher in non-pCR vs pCR, negative sign if lower). For paired on-treatment vs baseline analysis, samples were analyzed using a moderated t-test with limma^37^ using a paired design.

For breast cancer cell lines, log_2_-transformed TPM (Transcripts Per Million) and shRNA dependency scores (D2_combined_gene_dep_scores.csv) were downloaded from the DepMap resource (https://depmap.org). TNBC subtype information for cell lines was retrieved from Lehmann et al^38^ and metabolic subtypes were retrieved from Gong et al^20^. Associations between *OXA1L* RNA with TNBC subtypes and metabolic subtypes were assessed by ANOVA and associations between *OXA1L* dependency scores with *OXA1L* RNA levels were assessed by Pearson correlation.

### MRP signature generation

The MRP signature comprises all MRPL and MRPS genes quantified in TNBC PDXs at the protein level. Signature scores for individual PDX and clinical samples were calculated using single sample Gene Set Enrichment Analysis (ssGSEA)^39^ implemented through the ssGSEA2.0 tool (https://github.com/broadinstitute/ssGSEA2.0) and reported as normalized enrichment scores (NES). NES values were then used to assess associations with therapeutic response.

### Cell Culture

The MDA-MB-231 and MCF10a cell lines were initially purchased from the American Type Culture Collection, while the SUM159pt cell line was initially purchased from BioIVT. MDA-MB-231 cells were cultured in RPMI-1640 glucose-free media (Gibco, 11879020) supplemented with 5 mM glucose (Gibco, A2494001), 10% heat-inactivated FBS (R&D, S11550), and 1% antibiotic-antimycotic solution (Corning, 30-004-CI). SUM159pt cells were cultured in the same base media as MDA-MB-231 cells and supplemented with the same glucose and Antibiotic-Antimycotic concentrations. SUM159pt cells were also supplemented with 5% heat-inactivated FBS (R&D, S11550), 5 mg/ml insulin (Santa Cruz Biotechnology, sc-360248), and 1 mg/ml hydrocortisone (Sigma-Aldrich, H4001). MCF10a cells were cultured in MEGM Mammary Epithelial Cell Growth Medium BulletKit (Lonza, CC-3150) except for the substitution of the GA-1000 with 1% antibiotic-antimycotic Solution (Corning, 30-004-CI). All cells used in the experiments were maintained at fewer than 20 passages.

### Measurement of cell viability and confluency

Cells were counted using ViaStain AOPI staining solution (Revvity Health Sciences Inc. CS2-0106), which was added to cells immediately prior to counting with an Nexcelom Cellometer (Revvity Health Sciences, Inc.). For ATP readouts, the CellTiter-Glo assay was used. Cells were measured using the Promega CellTiter-Glo^TM^ Luminescent Cell Viability Assay (PR-G7572) according to the manufacturer’s instructions. For cell count readouts, cells were stained with DAPI then counted using a Cytation5 (BioTek) fluorescent plate reader. Cell confluency was assessed longitudinally using an Incucyte S3-C2 (Sartorius). Cells were seeded in a 96-well plate (Corning #35072) and five live-cell images per well were taken with the 10x objective every four hours for the duration of the experiment. Cell confluency was analyzed using the Incucyte image analysis software (Sartorius). Relative cell viability was determined by staining cells with Hoechst (Invitrogen, 34580) and measuring fluorescence using an Agilent BioTek Synergy LX (model number: SLXFA-SN) with the blue cube (BioTek, part number: 1505006, EM 460/40).

### Authentication of cell lines and PDX models

Cell lines and all PDX models were tested quarterly for mycoplasma using the Universal Mycoplasma Detection Kit (ATCC, 30-1012K). Identity of cell lines and PDX models was validated by short-tandem repeat (STR) DNA fingerprinting at the M.D. Anderson Cancer Center Cytogenetics and Cell Authentication Core. For STR DNA fingerprinting analysis, the Promega 16 High Sensitivity Kit (Catalog #DC2100) was used, and the results were compared with online databases (DSMZ/ATCC/JCRB/RIKEN).

### Gene silencing

Transient siRNA-mediated gene knockdowns were conducted with 5 nM siRNA targeting human *OXA1L* (5’-CAGAAAAAACAUGGUAUUAAA-3’, 5’-CUGGAAUUUAUGCAUGUUGAU-3’, 5’- AAGUUUUCCAGUCGAAUCAGA-3’) and Mission siRNA universal negative control #1 (Sigma, SIC001-10NMOL). Knockdown experiments were conducted using 10cm and 15cm plates, depending on the amount of material required for the downstream analyses. For 10cm plates and 48-hour cell pellet harvests, cells were seeded at 300K, while for 120-hour cell pellet harvests, cells were seeded at 25K. For 15 cm plates and 48-hour harvests, cells were seeded at 750K, and for 120-hour harvests, cells were seeded at 75K.

For 10cm plates 48hr harvests, 7.5 µl of 10 µM siRNA was combined with 30 µL of Lipofectamine RNAiMAX (Invitrogen, 13778150) with 3 mL OPTI-MEM media (Gibco, 11058021) was prepared. Following 15 minutes of incubation, complexes were added to cells that had been seeded 48-hour before at densities above. For 10cm plates with 120-hour harvests, the amount of Lipofectamine RNAiMAX was adjusted to account for the smaller number of cells at the time of transfection, instead of 30 μL, 7.5 μL of Lipofectamine was used for cells that would be harvested 120 hours post-transfection. For the 15cm dish, lipofection reagents were scaled up 2.5 times for the respective endpoint harvest time. Cells were harvested 48-120 hours post-transfection either on ice using a cell scraper or with trypsin if cell counting was needed for downstream analyses.

### *In vitro* drug treatments

Chemotherapies were prepared as 10 mM stock solutions, aliquoted and stored at -80°C. Docetaxel (Selleck Chemicals S1148) was dissolved in DMSO and Carboplatin (Selleck Chemicals S1215) was dissolved in complete media. Aliquots of these stock solutions were thawed at time of use, then discarded. Cells were seeded two days prior to treatments such that, at the time of treatment, cells were ∼30% confluent. Docetaxel was administered at 1 - 5 nM and carboplatin 25 – 100 µM. Cells were collected 48-120 hours following treatment. Tigecycline (TIG; Selleck Chemicals S1403) and chloramphenicol (Selleck Chemicals S1677) were prepared fresh in DMSO. TIG and chloramphenicol doses are indicated within each figure legend. For combination treatments, drugs were administered simultaneously, and doses are indicated within each figure legend.

### DNA extraction and relative mtDNA to nDNA quantification

Cell pellets were snap-frozen in liquid nitrogen and stored at -80 °C. DNA extraction was conducted using the Mag-Bind® Blood and Tissue DNA HDQ 96 Kit (OmegaTek M6399-00) as per the manufacturer’s protocol. The optional steps in the manufacturer’s protocol: 10 μL of RNase A was added per sample, and the elution buffer was heated to 70 °C prior to elution, were taken. The extracted DNA concentration was determined using the NanoDrop 2000 (Thermo Scientific).

To quantify the relative amount of mtDNA per nDNA, qPCR primers for mtDNA^10^ were used, while nuclear DNA primers were used to quantify nDNA. The mtDNA primers were MT-ND1 (forward: 5′-ATGGCCAACCTCCTACTCCT-3′, reverse: 5′-TAGATGTGGCGGGTTTTAGG-3′), and MT-ND6 (forward: 5′-TGGGGTTAGCGATGGAGGTAGG, reverse: 5’-AATAGGATCCTCCCGAATCAAC-3′). The nDNA primers were RGPD1 (forward 5′- GTGGAGCCACTGAGAATGGT-3′, reverse: 5′-GCATGCCTGGCTGATTTTAT-3′), and FUNDC2P2 (forward 5′- TGAGTCAGTGGACCTTGCAG-3′, reverse: 5′-CAGAATGGTTTGCAAGCTGA-3′). qPCR was conducted using Universal SYBR Green Supermix (Bio-Rad, 1725121), 2-4 ng of extracted DNA, and 1 μM of primers. The qPCR conditions are as follows: 50 °C for 2min, then 95 °C hot start for 10min, followed by 40 cycles of 95 °C for 15 seconds and 60 °C for 1 minute. Relative copy number was then computed using the relative 2^-ΔΔCq^ method.

### RNA extraction and reverse transcription quantitative PCR (RT-qPCR)

Total RNA was extracted using TRIzol reagent (Invitrogen™, 15-596-018) according to the manufacturer’s instructions. Briefly, approximately 0.5 × 10⁵ cells were collected in TRIzol reagent. RNA was precipitated with isopropanol and washed with 75% ethanol. After resuspension in nuclease-free water at 55 °C for 10 min, RNA was treated with DNase using the TURBO DNA-free™ Kit (Invitrogen™, AM1907) according to the manufacturer’s instructions. RNA was further purified using the RNA Clean & Concentrator kit (Zymo Research, R1017). cDNA was synthesized using the SuperScript™ III First-Strand Synthesis System (Invitrogen™, 18080051). Quantitative PCR was performed using TaqMan™ Gene Expression Assays (Applied Biosystems™, 4369016) with FAM- or VIC-labeled probes for individual genes (Applied Biosystems™, 4448490). Real-time qPCR was carried out on a CFX Opus Real-Time PCR System (Bio-Rad) using CFX Maestro software. Relative gene expression was calculated using the 2^⁻ΔΔCq^ method. ΔCq was defined as Cq(target) − Cq(*TUBB*). ΔΔCq was calculated by subtracting the mean ΔCq of the control group (vehicle or siNT) from the ΔCq of each experimental sample (TIG treatment or siOXA1L).

### Western blotting

Cell pellets were lysed in RIPA buffer (Thermo Scientific, 89901) containing protease inhibitor cocktail (Roche, 11836153001) and phosphatase inhibitor cocktail (PhosSTOP, Roche, 4906837001). Proteins were quantified with Pierce^TM^ BCA Protein Assay Kits (ThermoFisher Scientific 23225). Criterion TGX precast Midi Protein Gels (4-20%, Bio-Rad, 5671095) were used for SDS-PAGE. Gels were transferred to nitrocellulose membranes (Bio-Rad 1620112) using a Trans-Blot Turbo Transfer System set at 2.5A, 25V, 7min (Bio-Rad, 1704150). EveryBlot Blocking Buffer (Bio-Rad, 12010020) was used to block membranes for 10 minutes at room temperature and for the dilution of antibodies. The following primary antibodies were used: anti-OXA1L (1:1,000, Proteintech, 21055-1-AP), anti-beta-tubulin (1:1,000, Cell Signaling Technology, CST2146), anti-beta-actin (1:1,000, Cell Signaling Technology, CST3700), anti-Phospho Histone H2AX (Ser139) (1:1,000, Cell Signaling Technology, CST2577), anti-Tom70 (1:3,000, Proteintech, 14528-1-AP), anti-Mito Profile total OXPHOS Human WB antibody cocktail (1:500, abcam, ab110411), anti-Complex I NDUFB8 (1:1,000, abcam, ab110242), anti-Complex I MT-ND3 (1:500, abcam, ab192306), anti-Complex II SDHB (1:1,000, abcam, ab14714), anti-Complex III UQCRC2 (1:4,000, abcam, ab14745), anti-Complex IV MT-COXI (1:2,000, Abcam, ab14705), anti-Complex IV MT-COXII (1:1,000, abcam, ab110258), anti-Complex IV MT-COXIV (1:1,000, Cell Signaling Technology, CST4844), anti-Complex V ATP5A (1:1,000, abcam, ab14748), anti-Complex V MT-ATP8 (1:1,000, Cell Signaling Technology, CST96857), anti-caspase-3 (1:1,000, Cell Signaling Technology), and anti-cleaved caspase 3 (Asp175) (5A1E) (1:1,000, Cell Signaling Technology).

Secondary antibodies were used at a dilution of 1:10,000 in blocking buffer: horseradish peroxidase-conjugated anti-rabbit IgG H&L (Abcam, ab205718) and horseradish peroxidase-conjugated anti-mouse IgG H&L (Abcam, ab205719). Membranes were incubated in secondary antibodies for one hour, with rocking, at room temperature. Membranes were then washed with PBS including 0.1% Tween-20 (PBST) three times each for 10 minutes. For detection, Clarity Western ECL substrate (Bio-Rad, 1705060) was used, and images were captured on a ChemiDoc Touch imaging system (Bio-Rad).

### Mitochondrial fractionation

Mitochondrial fractions were extracted by differential centrifugation, as previously described^40^. Briefly, cell pellets containing >5 million cells were resuspended in RSB hypo buffer (10 mM NaCl, 1.5 mM MgCl_2_, 10 mM Tris-HCl, pH 7.5) on ice to allow for swelling of the cells. Cell lysis was facilitated on ice by dounce homogenization via 50 strokes with the small clearance pestle (DWK, 885302-0002) in a 2mL tissue grinding tube (DWK, 885303-0002). Lysis was halted by the addition of MSH buffer (210 mM mannitol; 70 mM sucrose; 5 mM Tris-HCl; and 1 mM EDTA, pH 7.5). RSB and MSH buffers were supplemented with cOmplete Protease Inhibitor Cocktail (Sigma, 11836170001) and PhosStop-phosphatase inhibitor tablets (Roche, 04906837001) to inhibit phosphatase and protease activity, respectively, and all centrifugation steps were performed in a prechilled (4°C) centrifuge. Cell suspensions were centrifuged at 600 × *g* for 5 minutes, and the supernatant was transferred to a new tube by pipetting. Supernatants were then centrifuged at 1,200 × *g* for 5 minutes to pellet any remaining nuclei from the sample. The supernatants were again collected, then centrifuged at 7,000 × *g* for 10 minutes to pellet the mitochondria-enriched fraction. Supernatants containing cytosolic fractions were collected in fresh tubes. Cytoplasmic fractions were further processed by centrifuging at 16,000 × *g* for 20 minutes to pellet cellular debris. The supernatant (cytosolic protein) was then transferred to a new tube and stored at -80 °C.

The mitochondria-enriched pellet was resuspended in MSH buffer and spun at 7,000 × *g* for 10 minutes to wash mitochondria. After discarding the wash supernatant, the pellet was resuspended for its final wash in MSH buffer and spun at 10,000 × *g* for 10 minutes. The supernatant was discarded, and the crude mitochondrial pellet was resuspended in 50-100 μl MSH buffer and proteins were quantified using the Pierce BCA Protein Assay Kit (Thermo-Fisher, 23223), then stored at -80 °C.

### Blue native polyacrylamide gel electrophoresis (BN-PAGE)

25-60 μg of mitochondria in MSH suspension from the fractionation were aliquoted and pelleted at 10,000 × *g* for 10 minutes at 4°C. After removing any remaining supernatant and the MSH buffer, mitochondrial pellets were resuspended in 1X NativePAGE Sample Buffer (Invitrogen) with 1% digitonin (8 g/g; Invitrogen) by gentle pipetting (∼20 times). Samples were incubated on ice for 15 minutes to solubilize the mitochondrial proteins. The samples were then centrifuged in a precooled centrifuge (4 °C) for 30 minutes at 18,000 × *g* and supernatants were transferred to new tubes. Next, 1-2 μL of 0.5% G-250 Coomassie dye (Invitrogen) was added to the samples. Samples were then loaded into pre-cast NativePAGE Novex 3-12% Bis–Tris gels in the XCell SureLock Mini-Cell system (Invitrogen) according to the manufacturer’s protocol. Briefly, the entire apparatus was assembled and placed in an ice bath. The wells of the gel were filled with dark blue cathode buffer (1X NativePAGE running buffer and 1X cathode additive), then samples and NativeMark unstained protein standard (Invitrogen) were loaded. The inner chamber was filled with dark blue cathode buffer, and the outer chamber was filled with anode buffer (1X NativePAGE running buffer) until it was 1/3 full. The gel was run at 100V for 30 minutes followed by 150V for another 30 minutes. The dark blue cathode buffer was then replaced with the light blue cathode buffer (1X NativePAGE running buffer and 0.1X Cathode Additive), and the gel was run for an additional 90-120 minutes at 250 V.

### Imaging Coomassie-stained BN-PAGE gels

The gel was transferred into a container with 4°C DI water, quickly rinsed with fresh 4°C DI water, then incubated in Fix/Destain solution (40% methanol + 7% acetic acid) for 1 hour on an orbital shaker at room temperature. Halfway through the incubation, the solution was refreshed. After incubation, Fix/Destain solution was removed and replaced with DI water. The gel was incubated in DI water overnight on an orbital shaker at room temperature. White-light images were taken on ChemiDoc Touch imaging system (Bio-Rad).

### Immunoblotting of BN-PAGE gels

Proteins from BN-PAGE gels were transferred to a PVDF membrane using 1X NuPAGE Transfer Buffer (Invitrogen) and the Trans-blot Turbo transfer system (Bio-Rad) run at 2.5 amps for 18 minutes. Membranes were rinsed in DI water then incubated in 8% acetic acid for 15 minutes to fix the proteins. Membranes were then washed with DI water and allowed to dry air for ∼30 minutes or until dry. Once dry, membranes were rehydrated with 100% methanol, which also removed the blue Coomassie staining, then washed in DI water before blocking with EveryBlot blocking buffer (Bio-Rad) for 10 min. Membranes were then incubated at 4°C overnight in primary antibody. After washing with PBST three times for ten minutes, HRP-linked secondary antibody was added at room temperature for one hour. Preparation of primary and secondary antibodies and their dilutions were done the same way as for western Blotting. Membranes were subjected to three more ten-minute PBST washes, then developed using Clarity Western ECL substrate (Bio-Rad). Images were taken on ChemiDoc Touch imaging system (Bio-Rad).

### Seahorse analysis of mitochondrial function

Mitochondrial function was assessed using the Agilent Technologies XF Cell Mito Stress Test (Agilent Technologies, 103015-100) and the Seahorse XFe96 extracellular flux analyzer. Briefly, on the day of the assay, viable cells were counted by ViaStain AOPI staining solution (Revvity Health Sciences Inc. CS2-0106) and seeded at a density of 25,000 to 50,000 per well in 180 µl of RPMI seahorse assay media (Agilent Technologies 103576-100) on XFe96 cell culture microplates (Agilent Technologies, 103794-100) that had been previously treated with Poly-D-lysine hydrobromide (MedChem Express Cat # HY-139201A). Cells were then incubated in a CO2-free incubator at 37°C for one hour, and the calibrant cartridge was put into the Seahorse XFe96 extracellular flux analyzer for calibration. Next, the cell plate was put into the analyzer to run the assay. For the assay, mitochondrial inhibitory drugs were prepared as per the manufacturer’s guidelines in RPMI Seahorse Assay Media; oligomycin (1.5 µM), FCCP (1 µM), and rotenone A/antimycin (0.5 µM).

### Clonogenicity measurement

Cells were treated in 10 cm dishes with TIG at 50 µM or at the IC_50_ for each respective cell line (MDA-MB-231 8 µM or SUM159 3 µM). 48-hour after treatment, cells were counted using ViaStain AOPI staining solution (Revvity Health Sciences Inc. CS2-0106). Then a 10,000 cells per 10 mL cell dilution was made and either 100 or 50 cells per condition were plated in 6-well plates in triplicate. The media was refreshed every two days. Seven days after the initial replating, cells were stained with crystal violet (Sigma-Aldrich HT90132), washed, and left to dry. Colonies were manually counted the following day.

### Mitochondrial fluorescent non-canonical amino acid tagging (Mito-FUNCAT)

The method was adapted from Yousefi et al, 2021^41^ and Malik et al, 2023^42^. Cells were plated on coverslips (Thorlabs CG15) placed in a 6-well dish (Corning 3516) that had been previously sterilized with 70% ethanol and treated with 1X Poly-D-lysine hydrobromide prepared by the company manual (MedChem Express HY-139201A). Chemotherapy treatments were conducted two days after seeding, then two days after treatment, cells were labeled, stained, and mounted.

To label the cells, the media was aspirated, and the cells were washed with PBS. Cells were then given customized methionine-free complete media (Thermo Fisher Scientific, removed methionine from RPMI A1451701) and respective translation inhibitors for one hour. To inhibit cytosolic translation, 100 mg/ml cycloheximide (Research Products International C81040-1.0) was used, and to inhibit mitochondrial translation, 50 µM TIG (Selleck Chemicals S1403) was used. For the negative control, both inhibitors were administered at the previously listed concentrations. Next, 500 mM HPG (click-iT metabolic labeling reagent C10186, Invitrogen) was added to the cells for 30 minutes. After labeling, cells were incubated on ice for two minutes in buffer A (dH_2_0 base, 10 mM HEPES, 10 nM NaCl, 5 mM KCl, 300 mM sucrose, and 0.015% (w/v) digitonin). Buffer A was then removed, and cells were incubated in buffer A.1 (dH_2_0 base, 10 mM HEPES, 10 nM NaCl, 5 mM KCl, and 300 mM sucrose) for 15 seconds. Cells were then fixed with 4% paraformaldehyde (Alfa Aesar 47392-9M) for 30 minutes, with gentle rocking at room temperature. After fixation, cells were washed with PBS for five minutes, followed by incubation in 100 mM NH_4_Cl in PBS for 15 minutes. This was followed by three washes each for five minutes in staining solution (PBS base, 5% BSA, 5% tryptone, and 0.1% Triton X-100). Cells were washed briefly with 3% BSA in PBS. Then, the incorporated methionine analog, HPG, was “clicked” to a fluorescent tag using the Click-iT cell reaction buffer kit (Thermo Scientific C10269) based on the manufacturer’s instructions including 3 μM Azide (AlexaFluor 594 Azide, Thermo Scientific A10270) for 30 minutes at room temperature. Afterwards, cells were washed with 3% BSA in PBS.

To stain the mitochondria, cells were incubated in primary CoxIV antibody (Abcam Cat# ab33985) for one hour at room temperature (1:200 in 3% BSA). After incubation, cells were washed three times in 3% BSA for five minutes each. Then the cells were incubated in secondary antibody, goat anti-mouse Alexa Fluor 488 (Jackson ImmunoResearch Laboratory, Cat# 115-547-003, 1:500 dilution in 3% BSA). After incubation, cells were washed with 3% BSA and stained with DAPI for five minutes at room temperature. Following this, the cells were washed with 3% BSA and then incubated for five minutes in PBS including 500 mM NaCl. Last, cells were washed with PBS three times, then mounted on slides (Fisher Scientific 12-550-15) with mounting media (Invitrogen Prolong Glass Antifade Mountant P36982). Slides were stored at room temperature and protected from light until imaging, which was completed within 14 days.

Imaging was conducted with a Zeiss LSM 880 with Airyscan confocal microscope using a 63X oil objective. The channels and powers of the lasers used for imaging were as follows: channel 405 at 2.0 W, channel 488 at 4.5 W, and channel 561 at 15.0 W. The acquisition mode frame size was 1912x1912, and the zoom was set at 2. Z-stacks were taken with slices at 0.16 μM intervals. Images were then processed using Fiji ImageJ.

### Mito-FUNCAT image analysis

Fluorescence image analysis was performed using skimage, numpy, and scipy libraries to quantify colocalization of mitochondrial translation. Multi-channel TIFF images were processed to extract HPG-Azide 494 (Channel 1, marking nascent protein synthesis), COXIV 488 (Channel 2, labeling mitochondria), and DAPI (Channel 3, identifying nuclear regions). To ensure consistent background removal, thresholding was first applied on raw images using an Otsu-derived intensity cutoff from the HPG-only control (positive signal). Pixels below this threshold were masked before subsequent processing. After thresholding, fluorescence intensities were normalized using the maximum signal intensity of the retained pixels to prevent artificial enhancement of low signal. Nuclear and cytosolic compartments were segmented using thresholding on the DAPI channel, defining the nucleus as DAPI-positive regions and the cytoplasm as the inverse. Colocalization analysis was conducted separately for nuclear and cytoplasmic compartments. Pearson’s correlation coefficient was calculated between Channels 1 and 2 to assess global signal correlation. Percent colocalization in the nucleus was defined as the proportion of overlapping positive pixels within the DAPI mask, normalized to the total DAPI-positive area. Percent colocalization in the cytoplasm was calculated as the proportion of overlapping signal outside the DAPI mask, normalized to the total cytoplasmic area where either Channel 1 or Channel 2 exhibited positive signal. Mitochondrial translation per cell was determined by combining the positive pixels from Channels 1 and 2 that overlap within the cytoplasm and then was normalized to cell count. The proportion of actively translating mitochondria was determined by the difference between the overlapping positive pixels from channel 2 positive pixels (COXIV) divided by channel 2 positive pixels in the cytoplasm.

### Animal Studies

Animal studies were conducted under approved BCM IACUC protocol AN-8243, which was in accordance with the National Institutes of Health *Guide for the Care and Use of Laboratory Animals*. Following the Association for Assessment and Accreditation of Laboratory Animal Care guidelines, mice were ethically euthanized.

For spheroid cultures, TNBC PDX tumor pieces were orthotopically implanted into Scid/Bg mice and left to grow until 800-1200 mm^3^. Tumors were collected, finely minced with scalpels (VWR 100499-580 and 100499-578), and processed for spheroid culture described below.

The PIM001-P PDX model was established and characterized at the University of Texas MD Anderson Cancer Center^9,43^. Briefly, cryo-preserved cell suspensions were thawed, ViaStain AOPI counted using a Cellometer K2 (Nexcelom Bioscience), and viable cells were resuspended in a 1:1 volume mixture of Matrigel (Corning 354234) and complete cell growth medium (Dulbecco’s modified Eagle’s medium: F12, Cytiva Hyclone, SH30023.01) supplemented with 5% FBS. A total volume of 20 μl of 400,000 cells were unilaterally injected into the fourth mammary fat pad of each female Scid/Bg [C.B-17/IcrHsd-PrkdcscidLystbg J] mouse (Envigo).

The chemotherapeutic docetaxel (DTX; Hospira, 0409-0201-10) was supplied as a solution containing polysorbate 80 NF, 4 mg anhydrous citric acid USP, 23% v/v dehydrated alcohol USP, q.s., with polyethylene glycol 300 NF at 20 mg/kg. DTX was administered by i.p. injection at a dose volume of 2 mL/kg. The chemotherapeutic carboplatin (CRB; TEVA Pharmaceuticals, 0703-4248-01) was supplied at 10 mg/ml in water for i.p. injection at 50 mg/kg at a dose volume of 5 mL/kg. The antibiotic TIG (Selleck Chemicals, S1403) was supplied as 200 mg aliquots, which were then dissolved at 25 mg/mL in 10% DMSO + 90% sterile water and sonicated. The i.p. injection was 100 mg/kg at a dose volume of 4 mL/kg. Three cycles of TIG were given, consisting of daily administration for 10 days, followed by a 3-day break.

### PDX *ex vivo* spheroid studies

Tumor chunks were digested in Dulbecco’s modified Eagle’s medium (DMEM):F12 (Cytiva HyClone, SH30023.01) supplemented with 5% FBS, 0.45% collagenase A (Roche, 1088793), 0.086% Hyaluronidase (Sigma-Aldrich, H3506), and 2% BSA (Sigma-Aldrich, A9418) for 4 hours at 37°C while rotating. After the initial digestion period, cells were centrifuged at 170 × g for 8 minutes at room temperature, resuspended in red blood cell (RBC) lysis buffer (Invitrogen 00-433-57), and incubated at room temperature for 3 minutes with periodic inversion of the tube. At the end of the incubation, the RBC buffer was inactivated with a matched volume of PDX epithelial cell growth medium (DMEM:F12 (Cytiva Hyclone, SH30023.01) supplemented with 5% FBS and 1X antibiotics). Cells were centrifuged at 120 × g for 5 minutes at room temperature, and the supernatant was removed. Cells were then resuspended in trypsin (Corning 25-053-CI) for 3 minutes at 37°C, inverted periodically. The trypsin was quenched with 2X volume of PDX media and spun down at 170 x g for 5 minutes at room temperature. The supernatant was aspirated and cells were resuspended in Dispase solution (Stemcell Technologies, NC9886504) supplemented with DNase I (1mg/ml, Stemcell Technologies, NC9007308) for 5 minutes at 37°C with periodic inverting. The dispase was inactivated with an equal volume of PDX media, and cells were collected by centrifugation at 170 x g for 5 minutes at room temperature. The supernatant was aspirated and the cell pellet was resuspended in PDX media and passed through a 70 µm filter followed by a 40 µm filter. The initial cell count was determined, and MACS separation was initiated to enrich for human PDX tumor cells by depleting mouse stromal cells. MACS separation (Miltenyi Biotec 130-104-694) was performed according to the manufacturer’s protocol. The final cell count of the enriched human cells was obtained, and 5,000 cells were plated per 50 µl in growth medium in 96-well low-attachment plates (Corning 3474). Cells were treated with TIG and CRB 24 hours post-seeding and CTG assays were conducted 48 hours post-treatment.

### Association of proteomic profiles with PDX *ex vivo* spheroid responses

Baseline PDX tumor proteomics data was retrieved from the Lei et al, 2025 manuscript^18^ and Petrosyan et al, 2022 article^34^. PDX models were grouped based on sensitivity to single-agent TIG response; models with TIG monotherapy viability at 150 µM with <50% were classified as Sensitive, and those ≥50% were classified as Resistant. Differential protein expression between TIG-sensitive (n=2: BCM-4272, BCM-7649) and TIG-resistant (n=3: BCM-3904, BCM-3887, BCM-5438) PDX models was assessed using a moderated t-test implemented in limma on log10-transformed protein intensity values. Proteins were ranked by signed -log10(p-value), calculated as -log10(p-value) multiplied by the sign of the log fold change from the limma contrast (Sensitive vs Resistant). Gene Set Enrichment Analysis (GSEA) was performed using WebGestaltR, the R tool for WebGestalt, against the Reactome pathway database. Pathways with a false discovery rate (FDR) < 0.15 were considered significantly enriched. The top 10 positively enriched (Sensitive) and top 10 negatively enriched (Resistant) pathways by normalized enrichment score (NES) were visualized as a bidirectional bar plot.

### Immunohistochemistry (IHC)

Harvested tumors were immediately sliced with scalpels and fixed in 10% formalin (Sigma HT501128-4L) at room temperature for 48-hours, followed by three 10 minute washes with PBS. Tissues were stored at 4 °C in 70% ethanol until processing the fixed tumors into paraffin blocks, from which 3 µm sections were cut. Routine hematoxylin and eosin (H&E; Epredia, 72711 and 71311) staining was conducted. IHC was conducted with antibodies against Ki67 (Dako Cat# M70240 at 1:200) and CC3 (Cell Signaling Technologies, CST9661 at 1:50) following antigen retrieval with 0.1M Tris-HCL 15 min at full pressure (above 90 °C) with a pressure cooker.

### Statistical analysis

Data shown are representative of at least three biological experiments. The bar graphs indicate the mean, and error bars represent the standard error of the mean (SEM). GraphPad Prism and R software, as indicated within the figure legends, were used for some statistical testing. Outliers were identified by the ROUT method using Prism and removed. For multiple comparisons, the post-hoc either the Dunnett’s test Tukey test were used after the one-way ANOVA. When data were not normally distributed, Mann-Whitney tests were used.

## RESULTS

### Mitochondrial translation and OXA1L are associated with TNBC chemotherapy resistance in human and PDX cohorts

Publicly available proteomic data from a Clinical Proteomic Tumor Analysis Consortium (CPTAC) TNBC study demonstrated oxphos was among the most significant Hallarmks50 GSEA pathways associated with resistance to DTX+CRB in tumors prior to therapeutic exposure^19^. In light of our prior findings that mitochondrial function is a heightened vulnerability of chemo-refractory TNBC^9,10^, we sought to extend analyses of those data to gain more in-depth insights into correlates of TNBC chemoresistance (Fig S1A-C). We found mitochondrial translation-related pathways were significantly enriched in pre-treatment tumors of patients who went on to harbor residual cancer burden (RCB) compared to those with a complete pathologic response (pCR; Fig 1A, S1A-B). A subset of those patients provided on-treatment biopsies three days after initiation of DTX+CRB. Again, mitochondrial translation-related pathways were among the significantly enriched pathways in matched on-treatment relative to pre-treatment tumors (Fig 1A, S1C).

**Fig 1.**
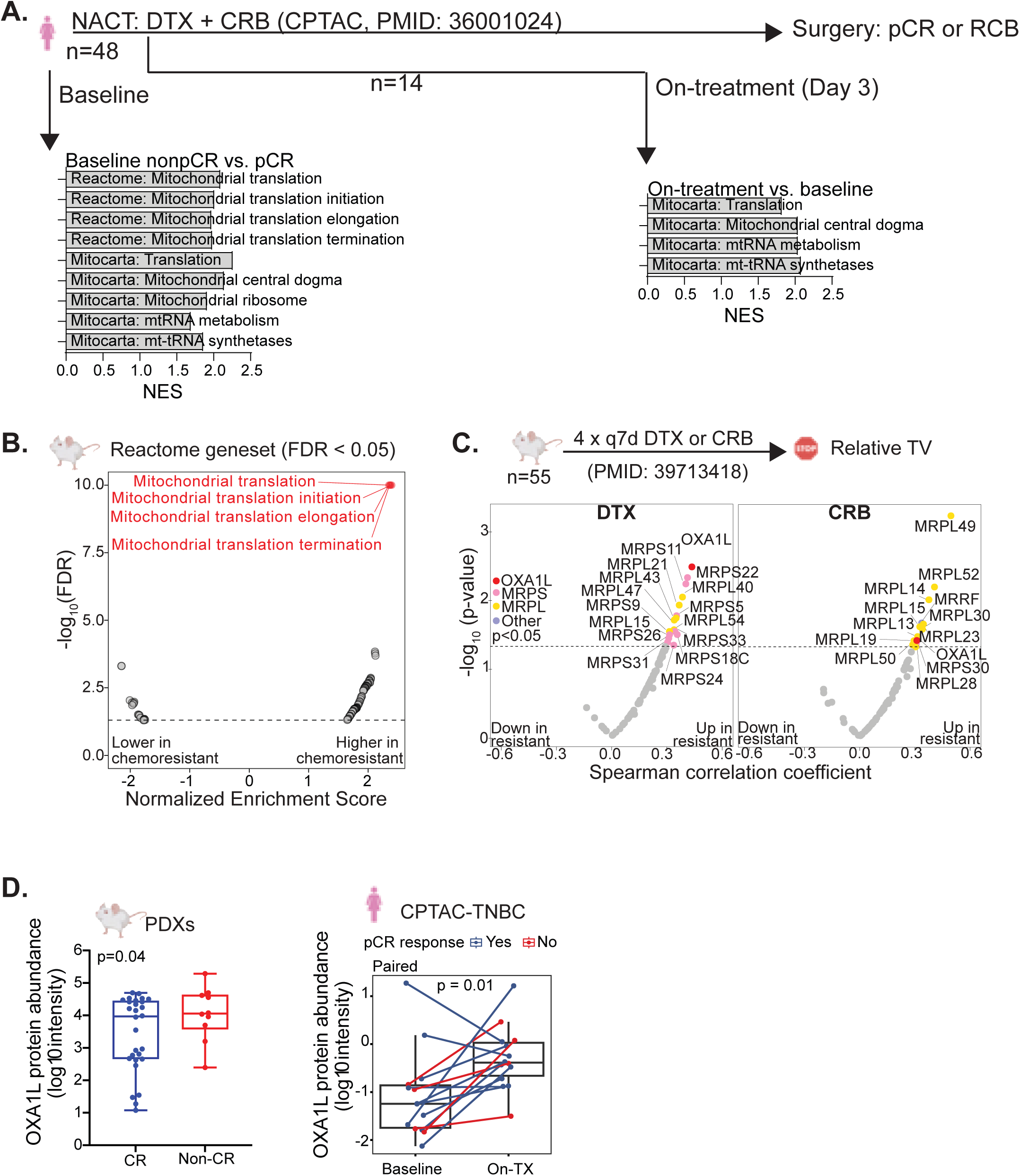
Mitochondrial translation signatures are enriched in chemoresistant TNBC and further induced by chemotherapy. (A) Mitochondrial-related pathways that were significantly enriched (FDR < 0.05) in non-pCR vs pCR CPTAC TNBC patient biopsies at baseline and on-treatment are shown. (B) GSEA Reactome pathway analysis of TNBC PDX models^18^, with only significantly altered pathways displayed. Pathways including the term ‘mitochondria’ are shown in red. (C) Leading edge proteins from the top pathway, ‘Reactome mitochondrial protein translation termination,’ upregulated in chemoresistant TNBC PDX models are shown^18^. Plotted are the -log_10_ p-value from Spearman correlations. The proteins above the dotted line are significantly enriched with p<0.05. (D) OXA1L protein abundance in TNBC patient^19^ and PDX^18^ cohorts comparing responsive and non-responsive tumors. P-values were derived from one-tailed t-tests and from limma moderated t-test with a paired design, respectively. Cartoon images were adapted from BioRender.

Complementing this, proteomic profiling of orthotopic TNBC patient-derived xenograft (PDX) tumors from a large-scale preclinical chemotherapy trial^18^ corroborated that prior to therapeutic exposure, the levels of proteins related to mitochondrial translation and ETC assembly were significantly higher in PDX models that were refractory to DTX and CRB than in those that were sensitive (Fig 1B). In fact, the Reactome ‘mitochondrial translation termination’ pathway was the top pathway associated with chemotherapy resistance among the 40 significantly upregulated pathways^18^ and was also enriched in the CPTAC dataset. Among the clinical and PDX datasets, there were 63 total leading-edge proteins in this pathway upregulated in chemoresistant samples (data not shown). Most proteins were mitochondrial ribosomal proteins (MRPs): 31 large subunit proteins (MRPLs) and 24 small subunit proteins (MRPSs), as well as mitoribosome recycling factor (MRRF), another component of the core mitoribosome machinery. The mitoribosome accessory protein OXA1L was, aside from core mitoribosome components, the protein most strongly associated with chemotherapy resistance that overlapped between the patient and PDX datasets (Fig. 1C-D). Additionally, OXA1L protein level significantly increased in the subset of the TNBC patient tumors biopsied on-chemotherapy treatment compared to matched pre-treatment tumors (Fig. 1D). We generated an MRP signature but did not observe a significant association with chemoresistance (Fig S1D). Given the dual functions of OXA1L in mitochondrial translation and insertion of ETC proteins into the IMM for ETC formation, these findings led us to hypothesize that mitoribosome function, largely supported by OXA1L, may be a major contributor to the metabolic rewiring^9,10^ that supports TNBC chemoresistance.

### OXA1L is required for ETC formation and mitochondrial function in human TNBC cells

Transient siRNA-mediated knockdown (KD) of *OXA1L* in MDA-MB-231 and SUM159pt human TNBC cells decreased the abundance of mtDNA- and nDNA-encoded ETC proteins comprising all five complexes, while not affecting cell confluency over time (Fig. 2A, S2A, & S3A). Notably, the reduction in protein levels became apparent five days after transfection, but not as early as two days, likely because the pool of existing ETC proteins had not yet turned over. The effect of *OXA1L* KD was specifically at the protein level, with no significant reduction in mtDNA or mtRNA levels observed in MDA-MB-231 and only a modest reduction in mtDNA with one siRNA in SUM159 cells (Fig. S3B-C). Concomitantly, oxygen consumption rate (OCR, *i.e.,* oxphos) significantly decreased, and extracellular acidification rate (ECAR, *i.e.* glycolysis) tended to non-significantly increase five days following *OXA1L* KD (Fig. 2B & S2B). ATP levels also significantly decreased upon KD (Fig 2C & S2C). Together, these findings indicate that *OXA1L* supports oxphos through protein-level regulation of mtDNA- and nDNA-encoded ETC proteins in human TNBC cells.

**Fig 2.**
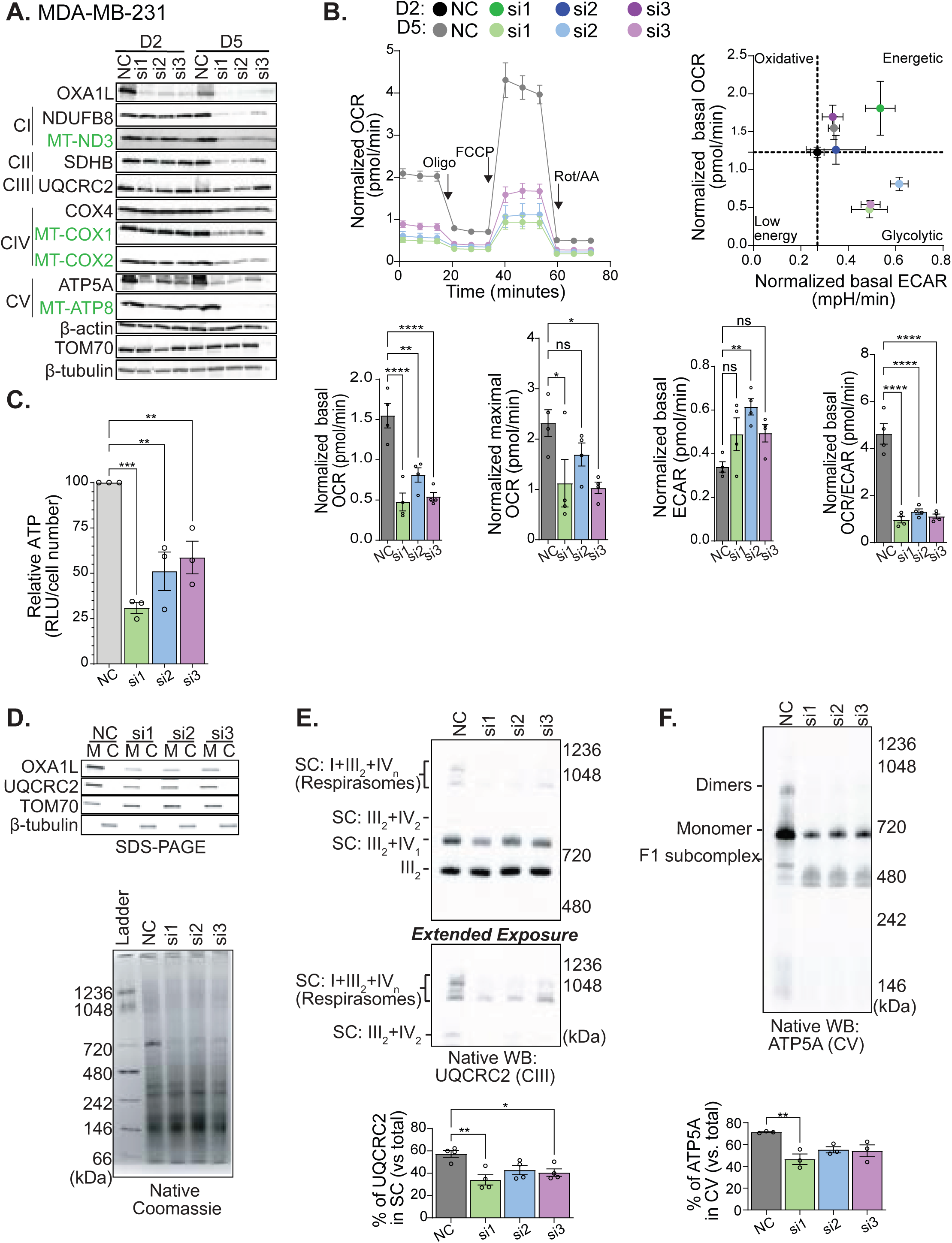
OXA1L is required for electron transport chain formation and function. MDA-MB-231 cells were transfected with siRNAs targeting *OXA1L* (si1, si2, si3) or a negative control sequence (NC). (A) Western blots of whole cell lysates probing for ETC proteins at day 2 (D2) and day 5 (D5) post-transfection are shown. (B) Seahorse MitoStress Tests of cells at D2 and D5 post-transfection. A representative biological replicate of the D5 timepoint is shown as the tombstone plot. All D2 and D5 biological replicates are represented in the energy plot. All biological replicates from D5 are shown in bar plots. OCR and ECAR levels were normalized to cell counts. (C) Relative ATP at D5 post-transfection. (D) Native Coomassie stain to assess SC formation with SDS-PAGE western blotting of mitochondrial and cytoplasmic fractions. (E) Blue native-PAGE immunoblotting for complex III: UQCRC2 and measurement of SC bands compared to total UQCRC2 quantified by densitometry. (F) Blue native-PAGE immunoblotting for complex V: ATP5A and measurement of monomer and dimer bands compared to total ATP5A quantified by densitometry. Blue native-PAGE immunoblotting for complex III using UQCRC2 and quantification of SC bands compared to total UQCRC2 with densitometry. Data are shown as mean ±SEM, and individual data points represent biological replicates. Significant comparisons are indicated; *p < 0.05, **p < 0.01, ***p < 0.001, ****p<0.0001 by one-way ANOVA.

Although *OXA1L* KD potently reduced the abundance of several key ETC proteins, some were only modestly decreased (Fig. 2A and S2A), possibly due to their inherent stability and turnover rates. This led us to ask how the formation of ETC complexes and supercomplexes (SCs) was affected by *OXA1L* loss. Blue native PAGE of proteins extracted from purified mitochondria, followed by Coomassie staining, showed an overall disruption of SC formation (Fig. 2D & S2D). Immunoblotting for UQCRC2 (CIII) revealed substantial disruption of partitioning into SCs, including respiring SCs (aka respirasomes), which have been reported to support oxphos efficiency^44,45^ (Fig. 2E & S2E). Furthermore, *OXA1L* KD dramatically impaired full assembly and dimerization of ATP synthase (Fig. 2F & S2F). Together, these findings reveal OXA1L is required for proper ETC protein levels, SC formation, and oxphos in human TNBC cells.

We used the Characterized Cell Line Encyclopedia gene dependency database results from RNAi screens^46^ to compare *OXA1L* dependence among breast cancer cell lines. Within TNBC, there was no significant correlation between *OXA1L* mRNA levels and the dependency score (Fig S4A-B), suggesting that mitochondrial translation dependence is influenced by additional factors. Further, there was no significant association between *OXA1L* mRNA level or *OXA1L* dependency score with transcriptomic^38^ (Fig. S4C-D) or metabolic^20^ (Fig. S4E) subtypes of TNBC (Fig S4A-E). Importantly, non-tumor immortalized mammary epithelial MCF10a cells had a lower *OXA1L* dependency score than all the TNBC cell lines included, except for the CAL148 cell line (Fig. S4A). KD of *OXA1L* in MCF10a cells had no significant effects on cell growth, ETC protein levels, or OCR^10^ (Fig. S4F-H). Together, these data suggest a selective dependency on OXA1L in TNBC, supporting a potential therapeutic window.

### OXA1L promotes oxphos adaptation and survival of chemotherapy refractory TNBC cells

OCR was significantly elevated by high-dose CRB (Fig 3A, S5A, S6A), consistent with our previous report of oxphos elevation by several DNA-damaging chemotherapies^10^. *OXA1L* KD abolished both the high- and low-dose CRB-induced OCR increases (Fig. 3A, S5A, S6A). Further, *OXA1L* KD potently reduced the abundance of mtDNA- or nDNA-encoded ETC proteins in the presence and absence of CRB (Fig. 3B & S5B). Moreover, *OXA1L* KD significantly increased sensitivity to CRB treatments (Fig 3C & S5C).

**Fig 3.**
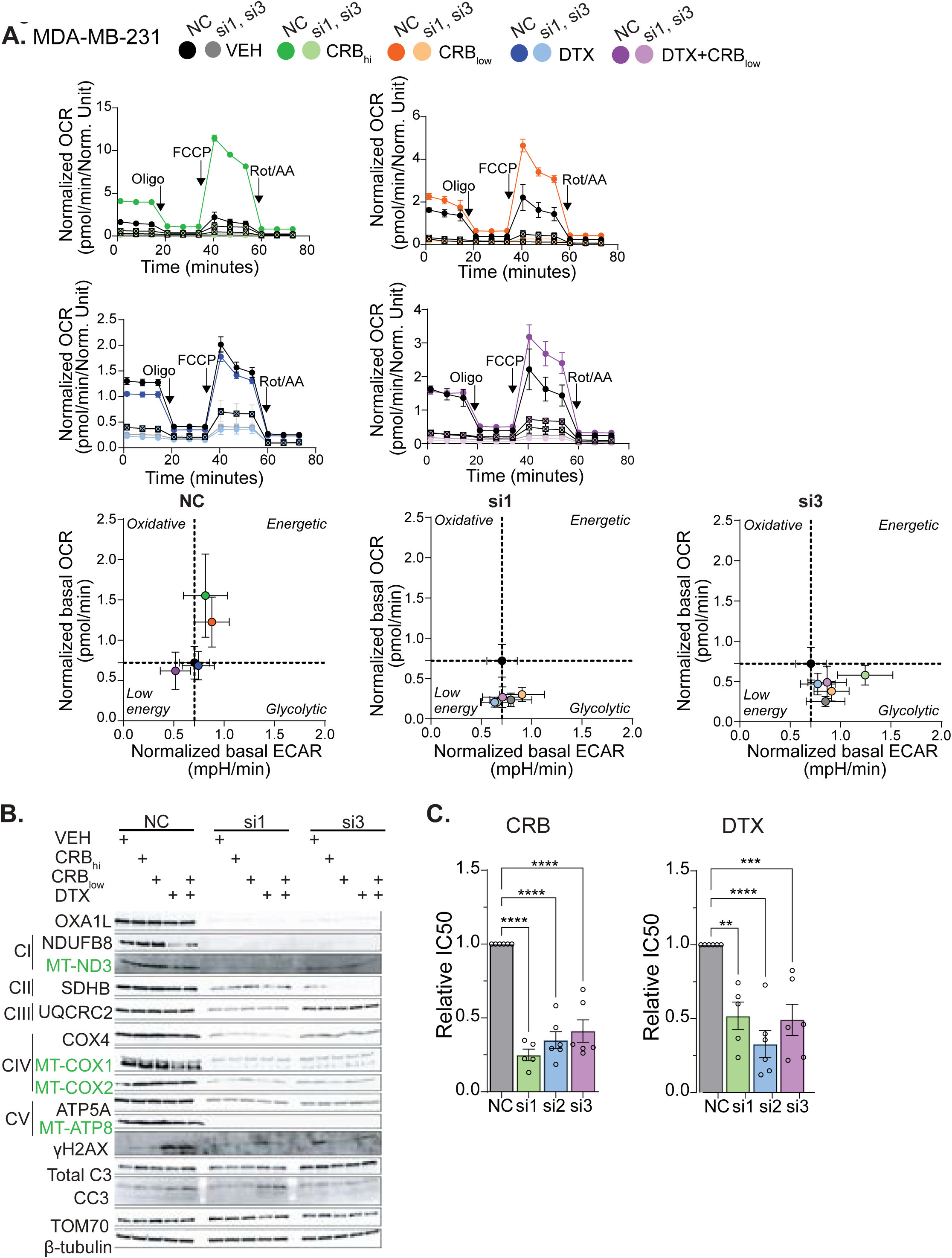
OXA1L is required for chemotherapy-induced metabolic adaptation and promotes TNBC resistance to chemotherapy. MDA-MB-231 cells were transfected with siRNAs targeting *OXA1L* (si1, si2, si3) or the negative control (NC) followed by treatment with chemotherapy. Chemotherapy treatments were vehicle (VEH), CRB_hi_ (100 µM), CRB_low_ (25 µM), DTX (5 nM), and DTX+CRB_low_. Readouts were conducted at D5 post-transfection and D2 post-chemotherapy treatment. (A) Seahorse MitoStress Tests of cells at D5 post-transfection. A representative biological replicate is shown in the tombstone plot, while all biological replicates are represented in the energy plot and bar graphs. OCR and ECAR values were normalized to viable cell counts. (B) Western blotting of whole cell lysates probing for ETC proteins. (C) IC_50_ values were measured in eight-point dose curves and relative values are displayed. Data are shown as mean ±SEM, and individual data points represent biological replicates. Significant comparisons are indicated; *p < 0.05, **p < 0.01, ***p < 0.001, ****p<0.0001 by one-way ANOVA.

Platinum chemotherapies are often combined with taxanes in the neoadjuvant setting for TNBC. Because OXA1L was identified as a leading edge protein associated with resistance to CRB as well as DTX^18^ in the PDX cohort, we also sought to test the contribution of OXA1L to DTX response. DTX caused a slight elevation of OCR in SUM159pt cells and no change in OCR in MDA-MB-231 cells. In both cases, *OXA1L* KD reduced oxphos when combined with DTX (Fig 3A, S5A, S6A). OXA1L significantly enhanced DTX sensitivity in both cell lines (Fig 3C & S5C). It is important to note that when DTX and CRB were combined, as is commonly used in the clinic, OCR was elevated, albeit to a lesser extent than with CRB alone (Fig. 3A, S5A, S6A). Then, with *OXA1L* KD, we observed a diminished OCR increase when DTX was combined with CRB. Together, these findings demonstrate the requirement for OXA1L for both oxphos and chemoresistance in chemo-refractory TNBC cells.

### Repurposed antibiotics disrupt mitochondrial translation and metabolic adaptations in TNBC cells

Several conventional antibiotics inhibit the bacterial ribosome, which shares high structural homology with the mammalian mitoribosome^47^ based on the theory of endosymbiosis^47^. Previous research suggests inhibiting mitochondrial translation using Tigecycline (TIG), an FDA-approved antibiotic used for treatment of sepsis, has anticancer activity in preclinical models of liver, breast, and ovarian cancer^47–51^, although the mechanistic relationship with mitochondrial translation remains unexplored. We hypothesized that combining TIG with conventional chemotherapies would disrupt the pro-survival mitochondrial adaptations elicited by chemotherapies. It is important to point out that, unlike OXA1L, TIG is only expected to disrupt mitoribosome activity and does not have a documented role in the insertion of nDNA-encoded ETC proteins into the IMM. We found the IC_50_ of TIG in human TNBC cells to be 3-8 mM by ATP-based or cell counting assays, which yielded highly concordant results (Fig. 4A & S7A-B), a six to 16-fold lower dose than used previously in breast cancer *in vitro* models to perturb cancer stem-like cell features^47^. We were able to recapitulate the previously observed potent inhibition of tumor initiation capacity with 50 µM TIG, but not with the IC_50_ doses (Fig. S8A-D). Interestingly, TIG treatment did not appear to increase apoptosis, with a notable reduction in caspase 3 cleavage compared to vehicle-treated cells (Fig. 4B, S7C), consistent with its seemingly cytostatic effects on TNBC cells quantified longitudinally (Fig. S8E).

**Fig 4.**
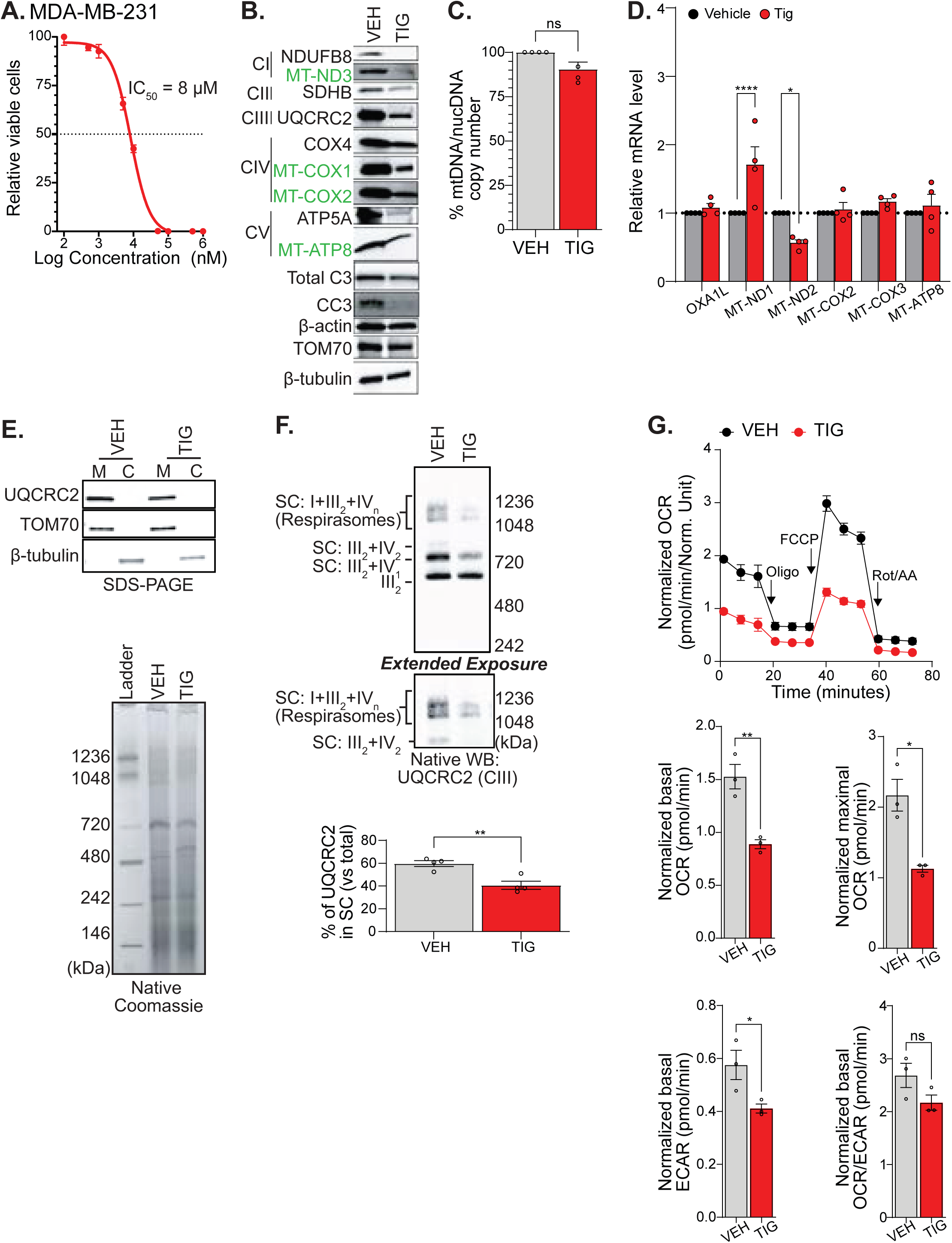
Mitochondrial function can be inhibited using repurposed antibiotics in TNBC. MDA-MB-231 cells were treated with vehicle (VEH) or TIG 8 µM for two days. (A) IC_50_ dilution curve, (B) Western blotting of whole cell lysates probing for ETC proteins, (C) mtDNA:nDNA ratio, and (D) qRT-PCR analysis of mitochondrial gene expression. Expression of mtDNA-encoded genes was normalized to *TUBB*. (E) Coomassie staining of a native gel to assess SC formation, and SDS-PAGE western blot of mitochondrial and cytoplasm fractions. (F) Blue native-PAGE immunoblotting for complex III: UQCRC2 and measurement of SC bands compared to total UQCRC2 by densitometry. (G) Seahorse MitoStress Tests showing a representative biological replicate for the tombstone plot and all biological replicates in the bar graphs. OCR and ECAR values were normalized to viable cell counts. Data are shown as mean ±SEM, and individual data points represent biological replicates. Significant comparisons are indicated; *p < 0.05, **p < 0.01, ***p < 0.001, ****p<0.0001 by unpaired t-test.

The abundance of ETC proteins (both mtDNA- and nDNA-encoded, comprising all five ETC complexes), but not mtDNA or mtRNA levels (except for *MT-ND1* in MDA-MB-231 cells), decreased with only 48 hours of TIG treatment (Fig. 4B-D & S7C-E) in both cell lines. Interestingly, we observed significant increases in several mtRNA transcript levels in SUM159pt cells upon TIG treatment, perhaps reflecting an adaptive response to translation inhibition. ETC respirasome formation was significantly decreased by TIG (Fig 4E-F & S7F-G), and we hypothesize this may have contributed to the reduction of nDNA-encoded ETC proteins induced by TIG. Indeed, there is evidence that SC assembly supports the stability of proteins within them^44^. Concordantly, TIG significantly reduced OCR (Fig. 4G, S7H).

To directly test the ability of TIG to inhibit mitochondrial translation, we conducted Mito-FUNCAT^41,42^, enabling quantification of nascent peptide levels based on labeling of the methionine analog HPG. Mitochondrial translation was measured by localizing HPG within mitochondria while inhibiting cytosolic translation with cycloheximide (CHX). As expected, CHX inhibited cytosolic translation while high-dose TIG did not (Fig S9A). Furthermore, one hour of treatment with high-dose TIG reduced mean mitochondrial translation per cell (number of overlapping pixels of HPG-azide with COXIV) by approximately 86% (Fig. S9A), while two days of TIG at its IC_50_ reduced mean mitochondrial translation per cell by approximately 36% (Fig. 5A). Two days of CRB significantly increased the proportion of mitochondrial area harboring active translation, whereas TIG significantly reduced this proportion as monotherapy or combined with CRB. Further, TIG significantly reduced total mitochondrial translation per cell when combined with either chemotherapy relative to the chemotherapy agent alone.

**Fig 5.**
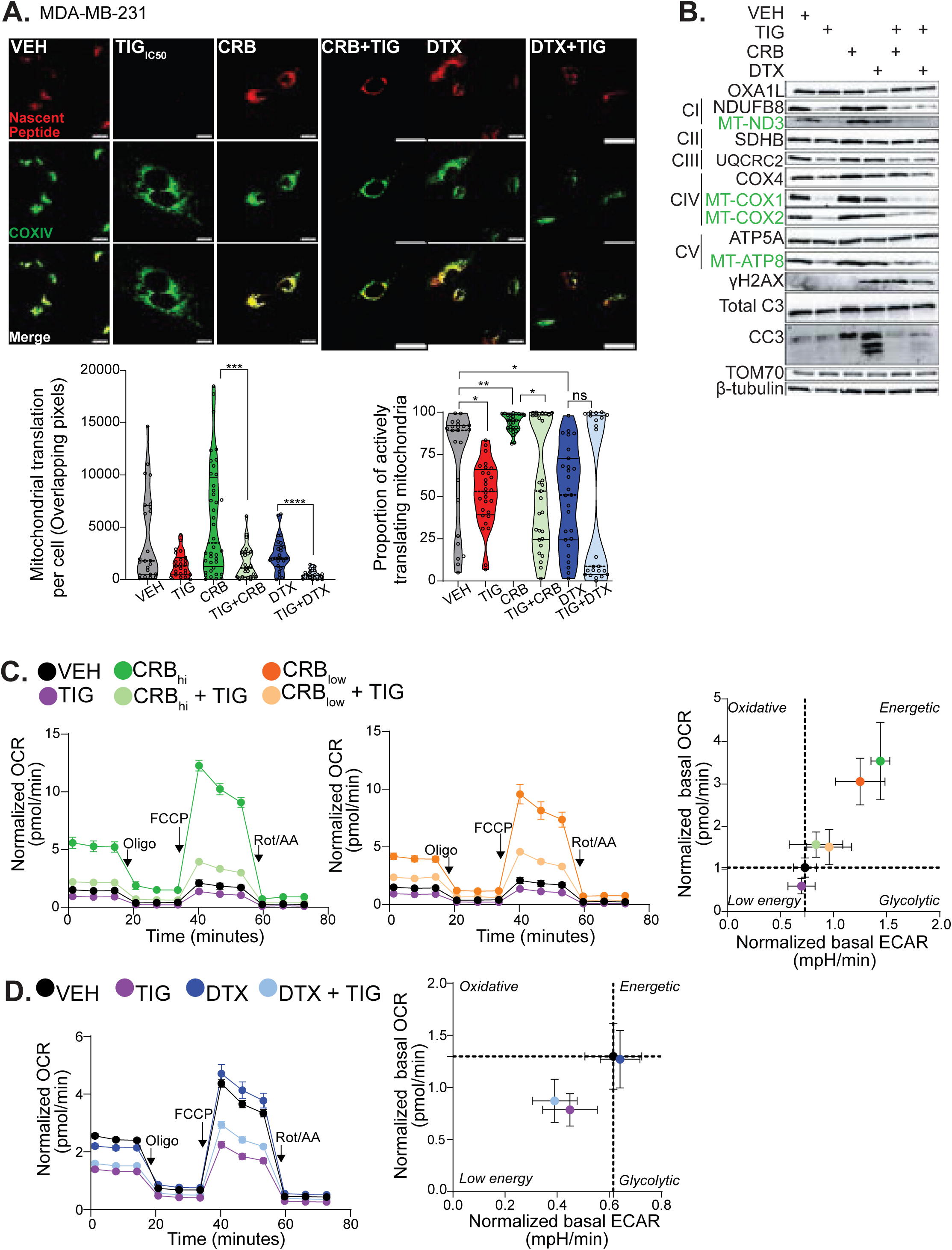
TIG suppresses chemotherapy-induced mitochondrial translation and metabolic adaptation. MDA-MB-231 cells were treated with VEH, TIG (8 µM), CRB (100 µM), and DTX (5 nM) for two days. (A) Representative images from Mito-FUNCAT are shown with nuclei masked (scale bar = 10 µmicron). The total colocalization of positive pixels for HPG-Azide and COXIV after nuclear masking per cell is shown in the first plot. In the second plot, the proportion of mitochondria actively translating is shown. Each point represents the quantification of an image. 25-30 cells were imaged per condition. Three independent biological replicates were conducted. The solid lines within the truncated violin plot are at the quartiles, and the dotted line is at the median. Normality and lognormality statistical tests revealed data were non-normal, so Mann-Whitney U Tests were performed between treatment pairs of interest. Significant comparisons are indicated; *p < 0.05, **p < 0.01, ***p < 0.001, ***p<0.0001). (B) Western blotting of whole cell lysates probing for ETC proteins. (C&D) Seahorse MitoStress Tests showing representative tombstone plots from one biological replicate. The energy plot includes all the biological replicates conducted. Data are shown as mean ±SEM, and individual data points represent biological replicates. Significant comparisons are indicated; *p < 0.05, **p < 0.01, ***p < 0.001, ****p<0.0001 by one-way ANOVA.

This imaging approach afforded us the ability to appreciate the heterogeneity of mitochondrial translation within and between cells. In the case of CRB, the total amount of mitochondrial translation per cell was highly heterogeneous, with some cells actively producing high levels of nascent mitochondrial peptides while other cells were seemingly silent. In converse, DTX treatment resulted in a relatively homogeneous amount of mitochondrial translation per cell, but highly heterogeneous proportions of actively translating mitochondria. In other words, while most DTX-treated cells had similar total amounts of mitochondrial translation, some cells had larger proportions of non-translating mitochondria than did others. These patterns may reflect differences in how mitochondrial translation is regulated across mitochondria within individual cells

Concordantly, TIG combined with chemotherapy decreased ETC protein abundance and disrupted CRB-induced OCR elevation (Fig. 5B-C, S9B, S10A-B). TIG, combined with DTX, reduced OCR and ETC protein levels (Fig. 5D, S9C, S10C). Together, these data support the notions that human TNBC cells surviving CRB harbor increased mitochondrial translation and mitochondrial function and that TIG rapidly and potently disrupts this effect.

### Mitoribosome inhibition enhances chemotherapy responses in a variety of TNBC models

TIG significantly increased the sensitivity of MDA-MB-231 and SUM159pt cells to CRB and DTX (Fig. 6-A-D). An additional FDA-approved antibiotic, chloramphenicol, could similarly disrupt oxphos and chemoresistance in the context of CRB (Fig. S11A-B). Furthermore, as observed with *OXA1L* KD in the TNBC cells, TIG reduced ETC protein levels but did not inhibit OCR of MCF10a cells (Fig. S12A-B). Additionally, only the highest dose of TIG tested moderately sensitized MCF10a cells to CRB, while all other doses left CRB sensitivity unaltered or even slightly reduced (Fig. S12C).

**Fig 6.**
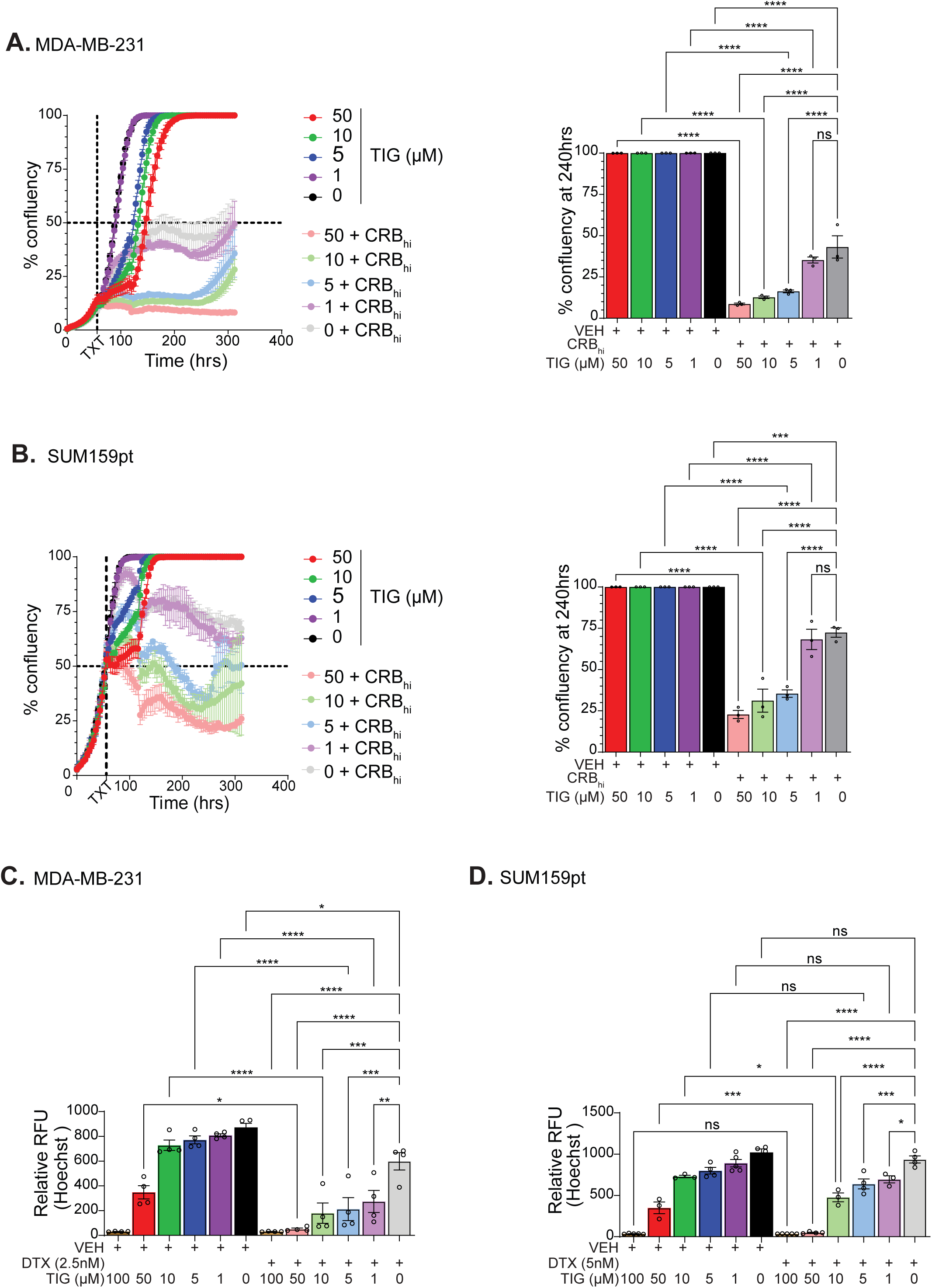
TIG sensitizes TNBC cells to chemotherapy. (A) Incucyte analysis of MDA-MB-231 cells treated with dilutions of TIG alone or simultaneously with CRB_hi_ (100 µM). The confluency of the cells readout at 240hrs (D10) was used for the one-way ANOVA followed by the post-hoc Dunnett’s test. (B) Incucyte analysis of SUM159pt cells treated with dilutions of TIG alone or simultaneously with CRB_hi_ (100 µM). The confluency of the cells readout at 240hrs (D10) was used for the one-way ANOVA followed by the post-hoc Dunnett’s test. (C) Hoechst measurement of relative cell counts on D7 of MDA-MB-231 cells after treatment with 2.5 nM of DTX followed by of TIG. (D) Hoechst measurement of relative cell counts on D7 of SUM159pt cells after treatment with DTX (5 nM) followed by TIG. Data are shown as mean ±SEM, and individual data points represent biological replicates. Significant comparisons are indicated; *p < 0.05, **p < 0.01, ***p < 0.001, ****p<0.0001 by one-way ANOVA.

Next, we tested the efficacy of TIG monotherapy and in combination with CRB in spheroid cultures from freshly digested tumors that had been depleted of mouse stroma from orthotopic PDX models, representing seven unique TNBC patients: PIM001-P^9,43^, BCM-4272, BCM-7649^52^, BCM-4195, BCM-3904, BCM-5438, and BCM-3887^52^. In all seven models tested combination of TIG with CRB had superior efficacy than CRB monotherapy (Fig. 7A).

**Fig 7.**
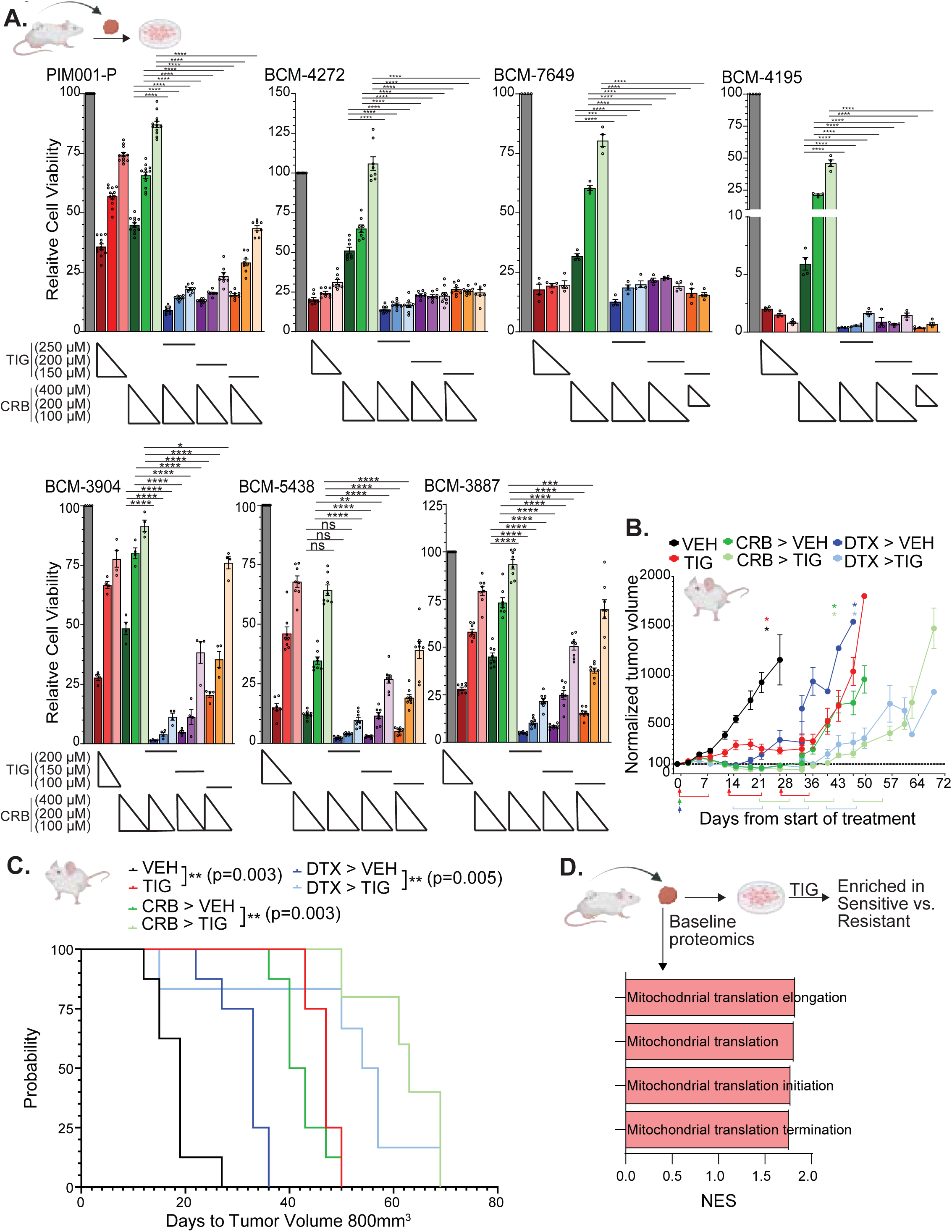
TIG chemo-sensitizes TNBC PDX *in vivo* and *ex vivo* models, with monotherapy efficacy correlating with mitochondrial translation pathway enrichment. Seven PDX models (PIM001-P, BCM-4272, BCM-7649, BCM-4195, BCM-3904, BCM-5438, and BCM-3887) were used to generate spheroid cultures for *ex-vivo* treatments with TIG and CRB. Two days following treatments, viability was measured by CTG luminescence. Significant comparisons are indicated; *p < 0.05, **p < 0.01, ***p < 0.001, ****p<0.0001 by one-way ANOVA followed by the post-hoc Tukey test. (B) The PIM001-P TNBC PDX model was orthotopically implanted into female SCID/bg mice to assess TIG given as a monotherapy or at the residual state following either DTX or CRB treatment. There were 8 mice per treatment group. Normalized tumor volumes are shown. Mice were treated upfront with one dose of CRB (dark green arrow), one dose of DTX (dark blue arrow), or a cycle of TIG (red brackets). In the CRB and DTX treatment arms, TIG treatment cycles (light green or blue brackets) began at the respective residual timepoint (the approximate tumor volume nadir) for each chemotherapy agent. Asterisks represent when mice on TIG treatment were collected. (C) Kaplan-Meier plot of days to reach 800 mm^3^ tumor volume. Log-rank p-values are displayed. (D) Mitochondria-related pathways from GSEA Reactome analysis that were enriched (FDR < 0.15) in TIG-sensitive PDX models compared to TIG-resistant PDX models based on spheroid treatment responses.

We then tested whether TIG could improve control of residual tumors persisting following ‘NACT’ in our previously characterized *in vivo* PDX model of residual TNBC, PIM001-P, in which tumor regrowth invariably follows initial regressions induced by a variety of conventional chemotherapies^9,53^. TIG monotherapy significantly slowed tumor growth (Fig. 7B-C). TIG treatment of residual tumors, defined as such when they reached their volume nadir after a single dose of CRB or DTX, significantly delayed their regrowth (Fig. 7B-C). TIG-treated tumors exhibited reduced levels of mtDNA-encoded ETC proteins (Fig. S13A) and Ki67 as monotherapy and when administered following chemotherapy. Additionally, TIG increased cleaved caspase-3 levels in combination-treated tumors (Fig. 7E). TIG treatment was associated with transient loss of 5-10% body weight that recovered during drug holidays, with no overt abnormalities observed in major organs (Fig. S13B). Despite TIG’s demonstrated clinical safety in its routine use, peritonitis of the mice’s abdominal tissues was visually apparent.

We next searched for proteomic correlates of TIG response to explore the future potential of patient stratification based on molecular profiles. Of the seven PDX-spheroid models tested (Fig, 7A), five had PDX mass spectrometry proteomics data available^18,34^. Exploratory Reactome pathway analysis revealed that the most TIG-sensitive models (those with TIG monotherapy viability at 150 µM that was <50% were classified as Sensitive) were enriched for pathways related to mitochondrial translation (Fig. 7D) as well as several additional cellular processes (Fig. S13D). OXA1L, along with many mitoribosome core components, was among the leading-edge proteins within those pathways (Fig. S13E). Notably, these mitochondrial translation-related pathways and leading-edge proteins substantially overlapped with those enriched in chemoresistant versus chemosensitive TNBC patients and PDXs (Fig. 1). Together, these findings demonstrate that TIG has monotherapy efficacy and enhances chemotherapy response in TNBC and identify a mitochondrial translation-related protein signature as a shared correlate of both chemotherapy resistance and TIG sensitivity, supporting its further evaluation as a putative predictive biomarker.

## DISCUSSION

In humans, the mitoribosome translates the 13 mtDNA-encoded transcripts in the mitochondrial matrix, all of which are components of ETC Complexes I, III, IV, and V. The remaining ETC components are encoded in nDNA. OXA1L has been studied most thoroughly in yeast, and its function in mammalian cells has been recently established. Its two reported roles are: 1) orchestrating translation termination of the 13 mtDNA-encoded ETC proteins^25,26^, and 2) co-translational insertion of mtDNA-encoded, as well as nDNA-encoded, ETC proteins into the IMM^24–27^. Additionally, OXA1L has been linked to mitochondrial diseases^27,29,32^, asthma^31,54^, osteoporosis^55,56^, pain^57^, diabetes complications^58^, liver disease^59^, Parkinson’s disease^33^, and cancer^60–63^. Despite these associations, there are limited functional studies of OXA1L in any disease, and no functional studies of OXA1L in the context of cancer have been reported. Our study provides evidence that OXA1L is crucial for mitochondrial translation, ETC formation and function, oxphos, and chemoresistance in TNBC. Its appearance across multiple external TNBC datasets suggests it’s potentially generalizable role in human TNBC.

*OXA1L* KD disrupted mitochondrial protein translation, ETC formation, and mitochondrial function. Those phenotypes became apparent five days, but not two days, after KD, likely because the existing pool of ETC proteins had yet to be turned over to uncover the effect^64^. Furthermore, we observed a decrease in both mtDNA-encoded and nDNA-encoded ETC proteins following KD, supporting the notion that OXA1L has dual functions in mitochondrial translation and the insertion of ETC proteins into the IMM. ETC complexes can stand alone or be in SCs with other complexes. In addition to other functions, SC formation has been shown to enhance electron flow efficiency and stabilize the individual ETC proteins comprising the complexes^44,45,65^. We observed a significant decrease in SC formation with *OXA1L* KD, which could explain the decrease in nDNA-encoded ETC protein levels.

It is important to note that we did not observe changes in mtDNA-encoded proteins or OCR in the immortalized, non-transformed mammary epithelial MCF10a cells with *OXA1L* KD. We previously reported that MCF10a has a higher mitochondrial copy number^10^, which could explain why these cells were less affected by the *OXA1L* perturbation than the TNBC cell lines. Further, chemotherapy treatments did not elicit mitochondrial rewiring in MCF10a cells^10^, suggesting nontumor cells may be more ‘resistant’ to mitochondrial perturbations than are tumor cells. This finding supports the notion that there may be a therapeutic window for translation of mitochondria-targeting therapies to the clinic.

OXA1L was required for the CRB-induced OCR boost in residual cells and increased the sensitivity of TNBC cells to both CRB and DTX *in vitro*. In our previous studies, we demonstrated that reducing mitochondrial function, induced by chemotherapy, made cells more sensitive to DNA-damaging agents^9,10^ so the increase in sensitivity to CRB upon *OXA1L* KD was somewhat expected. We speculate that the increased sensitivity to DTX upon *OXA1L* KD could be due to the cell’s limited “reservoir” of oxphos capacity, with KD diminishing it below a threshold required for cells to respond to the stress of chemotherapy. Another possibility is that DTX treatment induced greater ROS^10^, and when combined with OXA1L reduction, previously shown to induce ROS^32^, ROS accumulated to a lethal level. Future molecular studies will shed important light on these potentially divergent mechanisms.

Since there are currently no inhibitors of OXA1L, we focused on targeting mitochondrial translation using FDA-approved antibiotics, given the mitochondria’s bacterial ancestry. Treating TNBC cells at an IC_50_ dose of TIG disrupted mitochondrial protein translation, ETC formation, and mitochondrial function. In both MDA-MB-231 and SUM159pt cells, we observed an additive effect of combining TIG with chemotherapies, and we were able to recapitulate this finding with an additional antibiotic, chloramphenicol. Importantly, this additive effect was specific to the cancer cell lines tested, as it was not observed in MCF10a cells.

*Ex vivo* spheroid cultures derived from seven unique PDX models, five of which are annotated with mass spectrometry proteomic profiling data, afforded us the opportunity to conduct an unbiased search for correlates of TIG monotherapy sensitivity. This analysis revealed TIG efficacy is mechanistically rooted, with mitochondrial translation-related pathways, including OXA1L and many mitoribosome core proteins, enriched in sensitive models. Additionally, keratinization-related pathways were enriched in TIG sensitive models, although the functional relevance of this process is not yet clear. Interestingly, pathways related to epigenetic modifications and transcriptional regulation were enriched in TIG resistant models, perhaps suggesting compensatory mechanisms that may functionally support TIG resistance. Perhaps most importantly, the translation-related proteins and pathways associated with TIG sensitivity (Fig. 7D) were highly overlapping with those initially identified as associated with chemotherapeutic resistance (Fig. 1A-B). This suggests that targeting mitochondrial translation may be most fruitful in chemoresistant TNBCs. It is crucial these putative ‘biomarkers’ be validated in additional preclinical and clinical datasets.

Administration of TIG to CRB- or DTX-refractory residual tumors significantly delayed the regrowth of orthotopic PIM001-P PDX tumors. A limitation in the *in vivo* evaluation of TIG was its adverse effects on the abdominal cavity in mice, which had not been reported in previous mouse studies in the literature^48–51,66–71^. The dosage of TIG we administered to mice was 100 mg/kg, corresponding to a human equivalent dose^72^ of 8.13 mg/kg, that is about 12-fold less than human doses typically used to treat bacterial infections and about 37-fold lower than the clinical MTD determined in an AML cancer phase I clinical trial of TIG monotherapy (NCT01332786)^73^. Thus, clinically relevant doses of TIG were not achievable in the strain of mice we used with intraperitoneal administration. Additional animal strains and administration routes would be worth testing for further investigating the translational potential of TIG in TNBC. Our findings provide a foundation for future expanded studies to identify putative biomarkers of TIG sensitivity for clinical translation. Further, PDX mice lack a complete immune system, so future studies in immune competent TNBC models should be prioritized. It will also be important to consider the effects of antibiotics on the gut and breast microbiomes in the context of human TNBC, as microbiomes can influence tumor behaviors^74,75^.

## ACKNOWLEDGEMENTS

We are grateful to the breast cancer patients who donated their biopsies for cell line and PDX model generation. We are grateful to Janice Cowden and Amy Beumer for providing patient advocacy support for this research. Drs. Eric Jaehnig, PhD and Meenakshi Anurag, PhD aided in on-treatment CPTAC patient proteomic signature analysis. Dr. Junegoo Lee and Ms. Basmala Essam assisted with mouse experiments. Dr. Bora Lim provided clinical translation guidance. Dr. Helen Piwnica-Worms provided the PIM001-P PDX through a materials transfer agreement with the University of Texas MD Anderson Cancer Center, and model generation was supported by a generous gift from the Cazalot Foundation through the MD Anderson Women’s Cancer Moonshot Program.

BCM core facilities that supported this work included: The Optical Imaging & Vital Microscopy Core for Mito-FUNCAT imaging; The Mouse Metabolism and Phenotyping Core (NIH fund P30DK144025 & UM1HG006348) for Seahorse Bioanalyzer analysis; Patient-derived Xenograft Core (P30 Cancer Center Support Grant NCI CA125123, CPRIT Core Facilities Support Grant RP220646); Breast Center Pathology Core (supported by the Breast Center and a variety of research grants awarded to its faculty, including one of nine Specialized Programs of Research Excellence (SPORE) in Breast Cancer granted by the National Institute of Health. S10 grant (1S10OD028671-01) for digital imaging of the IHC slides) for processing tumor tissue into FFPE blocks and IHC staining. STR DNA fingerprinting was conducted by the Cytogenetics and Cell Authentication Core at M.D. Anderson Cancer Center.

## AUTHOR CONTRIBUTIONS

MJB and GVE were responsible for study conceptualization, carrying out the study, data presentation, and manuscript writing.

SWW conducted ETC complex analyses and provided experimental guidance under the supervision of GVE. MLB conducted qRT-PCR and provided experimental guidance under the supervision of GVE.

AL conducted mammary epithelial cell experiments and assisted with computational analyses under the supervision of GVE.

ASG conducted mouse experiments under the supervision of GVE.

KW conducted qPCR assays under the supervision of GVE.

LED aided in PDX experiments under the supervision of MTL.

IS conducted fluorescence colocalization quantitative analyses under the supervision of QZ.

QZ oversaw fluorescence colocalization quantitative analyses.

BZ aided in oversight of proteogenomic analyses.

JTL conducted computational analyses of transcriptomic and proteomic datasets under the supervision of BZ and MTL.

MTL aided in oversight of PDX studies and proteogenomic analyses.

## FUNDING

G.V.E. is a Cancer Prevention Research Institute of Texas (CPRIT) Scholar in Cancer Research. The authors were supported by CPRIT RR200009 to G.V.E.; National Institutes of Health (NIH) K22-CA241113 to GVE, R37CA269783-01A1 to G.V.E., and T32 predoctoral training grant T32GM136560-02 to MJB; an American Cancer Society Research Scholar Grant RSG-22-093-01-CCB to GVE; a Mary Kay Ash Foundation Cancer Research Grant 02-24 to GVE; a Breast Cancer Alliance Young Investigator Award to GVE; a National Science Foundation (NSF) Graduate Research Fellowship 2140736 to MJB; a Myra Branum Wilson Baylor Research Advocates for Student Scientists (BRASS) Scholarship to MJB; an American Cancer Society Postdoctoral Fellowship PF-24-1293970-01-TBI to SW; a Frank & Sandra Kimmel Endowment Postdoc Fellowship to MLB. The content is solely the responsibility of the authors and does not necessarily represent the official views of the NIH, NSF, CPRIT, ACS, or BRASS.

## SUPPLEMENTAL FIGURE LEGENDS

**Fig S1. Mitoribosome protein signatures are enriched in non-pCR clinical and non-responsive PDX TNBC tumors.** (A&B) Significant pathways enriched in the non-pCR vs pCR CPTAC TNBC patient biopsies at baseline from Gene Set Enrichment analysis with Reactome and MitoCarta geneset collections that was performed using WebGestalt^36^. Significance set at FDR < 0.05. (C) Significant pathways enriched in the on-treatment vs. baseline CPTAC TNBC patient biopsies from the Gene Set Enrichment analysis with MitoCarta geneset collections that was performed using WebGestalt^36^. Significance set at FDR < 0.05. (D) MRP protein signature comparing CPTAC-TNBC patient tumors^19^ and TNBC PDXs^18^ cohorts comparing responsive vs. non-responsive tumors. P-values derived from limma moderated t-test with a paired design and from t-test, respectively.

**Fig S2. OXA1L is required for ETC protein expression and mitochondrial function in SUM159pt cells.** SUM159pt cells were transfected with siRNAs targeting *OXA1L* (si1, si2, si3) or the negative control (NC). (A) Western blotting of whole cell lysates probing for ETC proteins at Day 2 (D2) and Day 5 (D5) post-transfection. (B) Seahorse MitoStress Tests of cells at D2 and D5 post-transfection. A representative biological replicate of D5 is shown as the tombstone plot. All D2 and D5 biological replicates are represented in the energy plot. All biological replicates from D5 are shown in the bar graphs. OCR and ECAR values were normalized to viable cell counts. (C) Relative ATP at D5 post-transfection. (D) Native Coomassie stain to assess SC formation with SDS-PAGE western blotting of mitochondrial and cytoplasmic fractions. (E) Blue native-PAGE immunoblotting for complex III: UQCRC2 and measurement of SC bands compared to total UQCRC2 quantified by densitometry. (F) Blue native-PAGE immunoblotting for complex V: ATP5A and measurement of monomer and dimer bands compared to total ATP5A quantified by densitometry. Data are shown as mean ±SEM, and individual data points represent biological replicates. Significant comparisons are indicated; *p < 0.05, **p < 0.01, ***p < 0.001, ****p<0.0001 by one-way ANOVA.

**Fig S3. *OXA1L* knockdown does not affect TNBC cell growth or mtDNA content**MDA-MB-231 and SUM159pt cells were transfected with siRNA targeting *OXA1L* (si1, si2, si3) or the negative control (NC). (A) Incucyte analysis of cellular confluence, (B) mtDNA:nDNA ratio quantification, and (C) qRT-PCR analysis of mitochondrial RNA levels. Expression of mtDNA-encoded genes was normalized to *TUBB*. Data are shown as mean ±SEM, and individual data points represent biological replicates. Significant comparisons are indicated; *p < 0.05, **p < 0.01, ***p < 0.001, ****p<0.0001 by one-way ANOVA.

**Fig S4. *OXA1L* knockdown does not significantly alter mitochondrial respiration in non-transformed mammary epithelial cells**. (A) *OXA1L* mRNA expression aligned with TNBC subtype, metabolic subtype (MPS), mRNA expression, and dependency score. The more negative the score is, the greater the gene dependency. (B) *OXA1L* dependency score plotted against *OXA1L* mRNA expression. P-values derived from Pearson Correlation. (C) TNBC subtype plotted against mRNA expression. (D) *OXA1L* dependency score plotted against TNBC subtype. (E) OXA1L dependency score plotted against MPS. (F) MCF10a cells were transfected with siRNAs targeting *OXA1L* (si1, si2, si3) or the negative control (NC). Incucyte analysis of cellular confluence with *OXA1L* KD. (G) Western blotting of MCF10a whole cell lysates on days D2 and D5 after transfection, probing for mtDNA-encoded proteins. (H) Seahorse MitoStress Tests of MCF10a cells D2 and D5 post-transfection. A representative biological replicate from D5 is shown in the tombstone plot. D2 and D5 timepoints from all the biological replicates are represented in the energy plot. Only D5 are shown in bar graphs. OCR and ECAR values were normalized to viable cell counts. Data are shown as mean ±SEM, and individual data points represent biological replicates. Significant comparisons are indicated; *p < 0.05, **p < 0.01, ***p < 0.001, ****p<0.0001 by one-way ANOVA.

**Fig S5. *OXA1L* KD impairs chemotherapy-induced metabolic adaptation and sensitizes TNBC cells to chemotherapy.** SUM159pt cells were transfected with siRNAs targeting *OXA1L* (si1, si2, si3) or the negative control (NC), followed by treatment with chemotherapy. Chemotherapy treatments were CRB_hi_ (100 µM), CRB_low_ (25 µM), DTX (5 nM), and DTX+CRB_low_. Readouts were conducted at D5 post-transfection and D2 post-chemotherapy treatment. (A) Seahorse MitoStress Tests of cells at D5 post-transfection. A representative biological replicate is shown in the tombstone plot, while all biological replicates are represented in the energy plot and bar graphs. OCR and ECAR values were normalized to viable cell counts. (B) Western blotting of whole cell lysates, probing for ETC proteins. (C) IC_50_ values were measured in eight-point dose curves, and relative values are displayed. Data are shown as mean ±SEM, and individual data points represent biological replicates. Significant comparisons are indicated; *p < 0.05, **p < 0.01, ***p < 0.001, ****p<0.0001 by one-way ANOVA.

**Fig S6. *OXA1L* KD suppresses CRB-induced increases in mitochondrial respiration.** MDA-MB-231 and SUM159pt cells were transfected with siRNAs targeting *OXA1L* (si1, si2, si3) or the negative control (NC). Following transfection, cells were treated with chemotherapy. OCR and ECAR values were normalized to viable cell counts. (A) Data from Seahorse MitoStress Tests are shown as mean ±SEM, and individual data points represent biological replicates. Significant comparisons are indicated; *p < 0.05, **p < 0.01, ***p < 0.001, ****p<0.0001 by one-way ANOVA.

**Fig S7. TIG inhibits mitochondrial translation in SUM159pt cells.** (A) Comparing ATP readout vs. cell count readout of MDA-MB-231 cells treated with TIG dilutions to determine IC50. SUM159pt cells were treated with vehicle (VEH) or TIG 3 µM for two days. (B) IC_50_ dilution curve, (C) western blotting of whole cell lysates probing for ETC proteins, (D) mtDNA:nDNA ratio, and (E) qRT-PCR analysis of mitochondrial gene expression. Expression of mtDNA-encoded genes was normalized to *TUBB*. (F) Coomassie staining of a native gel to assess SC formation, and SDS-PAGE western blot of mitochondrial and cytoplasm fractions. (G) Blue native-PAGE immunoblotting for complex III: UQCRC2 and measurement of SC bands compared to total UQCRC2 by densitometry. (H) Seahorse MitoStress Tests showing a representative biological replicate for the tombstone plot and all biological replicates in the bar graphs. OCR and ECAR values were normalized to viable cell counts. Data are shown as mean ±SEM from a minimum of three independent biological replicates. Significant comparisons are indicated; *p < 0.05, **p < 0.01, ***p < 0.001, ****p<0.0001 by unpaired t-test.

**Fig S8. TIG reduces TNBC cell viability in a dose-dependent manner.** MDA-MB-231 cells were treated with TIG at a high dose (50 µM) and the IC_50_ dose (8 µM). (A) Relative viable cells from AOPI counts on D2 after treatment. (B) Colony counts on D7 after seeding treated cells at 100 cells per well. SUM159 cells treated with TIG at a high dose (50 µM) and the IC_50_ dose (3 µM). (C) Relative viable cells from AO/PI counting on D2 after treatment. (D) Colony counts on D7 after seeding treated cells at 100 cells per well. (E) Incucyte analysis of MDA-MB-231 and SUM159pt cells treated with TIG dilutions. The confluency of the cells readout at the end of the experiment (120hrs) was used for the one-way ANOVA followed by the post-hoc Dunnett’s test. Data are shown as mean ±SEM, and individual data points represent biological replicates. Significant comparisons are indicated; *p < 0.05, **p < 0.01, ***p < 0.001, ****p<0.0001 by one-way ANOVA.

**Fig S9. TIG suppresses mitochondrial respiration in chemotherapy-treated TNBC cells.** (A) Mito-FUNCAT analysis of control samples of MDA-MB-231 cells. Cells were given HPG, a methionine analog, to be incorporated into nascent polypeptides, which were then fluorescently detected via click chemistry. Additionally, cells were treated with inhibitors to block either cytosolic translation (cycloheximide, CHX) or mitochondrial translation (TIG, high dose, 50 µM). Representative images from each condition are shown (scale bar = 10 µm), and the quantification of HPG-Azide and COXIV total colocalization of positive pixels for HPG-Azide and COXIV per cell after nuclear masking is plotted in the first plot. In the second plot, the proportion of mitochondria actively translating is shown. Each dot represents quantification of an image. 25-30 cells were imaged per condition. Three independent biological replicates were conducted. The solid lines within the truncated violin plot are at the quartiles, and the dotted line is at the median. (B) Seahorse MitoStress Tests of MDA-MB-231 cells treated with TIG (8 µM) as a monotherapy or in combination with CRB_hi_ (100 µM) or CRB_low_ (25 µM). (C) Seahorse MitoStress Tests of MDA-MB-231 cells treated with TIG (8 µM) as a monotherapy or with docetaxel (DTX 5 nM). Data are shown as mean ±SEM from a minimum of three independent biological replicates. *p < 0.05, **p < 0.01, ***p < 0.001, ****p<0.0001 by one-way ANOVA.

**Fig S10. TIG suppresses chemotherapy-induced metabolic adaptation in SUM159pt cells.** SUM159pt cells were treated with VEH, TIG (3 µM), CRB (100 µM), or DTX (5 nM) for two days. (A) Western blotting of whole cell lysates probing for ETC proteins. (B&C) Seahorse MitoStress Tests showing representative tombstone plots from one biological replicate. The energy plot includes all the biological replicates conducted. Data are shown as mean ±SEM, and individual data points represent biological replicates. Significant comparisons are indicated; *p < 0.05, **p < 0.01, ***p < 0.001, ****p<0.0001 by one-way ANOVA.

**Fig S11. Mitochondrial translation inhibition with chloramphenicol sensitizes TNBC cells to CRB.** MDA-MB-231 cells were treated with chloramphenicol. (A) Incucyte analysis of cells treated at different dilutions of chloramphenicol alone or with CRB_hi_ (100 µM). The confluency of the cells readout at the end of the experiment (164hrs) was used for the one-way ANOVA followed by the post-hoc Dunnett’s test. (B) Seahorse MitoStress Tests showing representative tombstone plots from one biological replicate. The energy plot includes all the biological replicates conducted. Data are shown as mean ±SEM, and individual data points represent biological replicates. Significant comparisons are indicated; *p < 0.05, **p < 0.01, ***p < 0.001, ****p<0.0001 by one-way ANOVA.

**Fig S12. TIG does not significantly alter mitochondrial function or chemotherapy sensitivity in non-transformed mammary epithelial cells.** MCF10a cells were treated with VEH, TIG (8 µM), or CRB (10 µM) for two days unless noted otherwise. (A) Western blotting of whole cell lysates probing for ETC proteins. (B) Seahorse MitoStress Tests showing representative tombstone plots from one biological replicate. The energy plot includes all the biological replicates conducted. (C) Incucyte analysis of cells treated with different dilutions of TIG alone or with CRB_hi_ (100 µM). The confluency of the cells readout at the end of the experiment (312hrs) was used for the one-way ANOVA followed by the post-hoc Dunnett’s test. Data are shown as mean ±SEM, and individual data points represent biological replicates. Significant comparisons are indicated; *p < 0.05, **p < 0.01, ***p < 0.001, ****p<0.0001 by one-way ANOVA.

**Fig S13. TIG tolerability and target engagement *in vivo* and the correlation of pathways and proteins with TIG response in PDX-derived spheroids.** Female Scid/bg mice bearing orthotopic PIM001-P tumors were treated with TIG and/or CRB. (A) Western blotting of tumors collected on treatment. (B) Normalized body weights to the start of treatment are plotted. Mice were treated upfront with one dose of CRB (dark green arrow), one dose of DTX (dark blue arrow), or a cycle of TIG (red brackets). In the CRB and DTX treatment arms, TIG treatment cycles (light green or blue brackets) began at the respective residual timepoint for each chemotherapy agent. Asterisks represent when mice on TIG treatment were collected. (C) Slides stained with H&E or antibodies against Ki67 and CC3 (scale bar = 50 µm) are shown from PIM001-P tumors collected on the last treatment day of the second cycle. Cartoons adapted from BioRender. (D) Pathway enrichment analysis of differentially expressed proteins between TIG-sensitive and TIG-resistant PDX models. Bar plots showing the top 10 Reactome pathways enriched in TIG-sensitive (red, right) and TIG-resistant (blue, left) models by Gene Set Enrichment Analysis (GSEA) of proteomic data (FDR < 0.15) are shown. The x-axis represents the normalized enrichment score (NES). (E) The top 20 leading edge proteins from Reactome mitochondrial translation pathways enriched in TIG-sensitive vs. resistant PDX-derived spheroids.

## Notes

### Summary of Updates

Since our original submission, this revised version has additional data that was recently generated and has been added to the manuscript.

