## Supplementary figures and images for "Mitochondrial translation is a targetable dependency of chemo-refractory triple negative breast cancer"

### Supplementary Figure 1

Supplementary Figure 1.

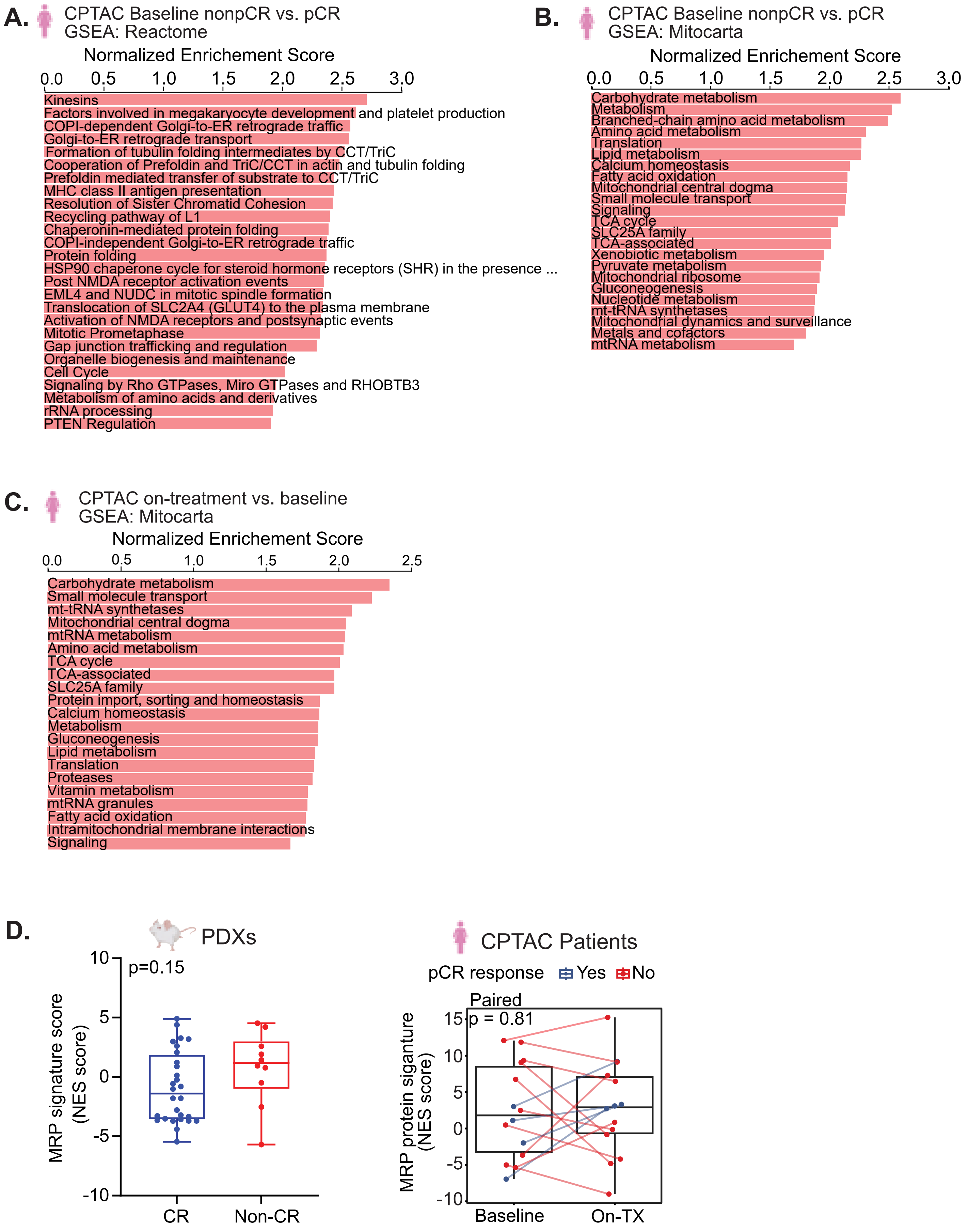

### Supplementary Figure 2

Supplementary Figure 2.

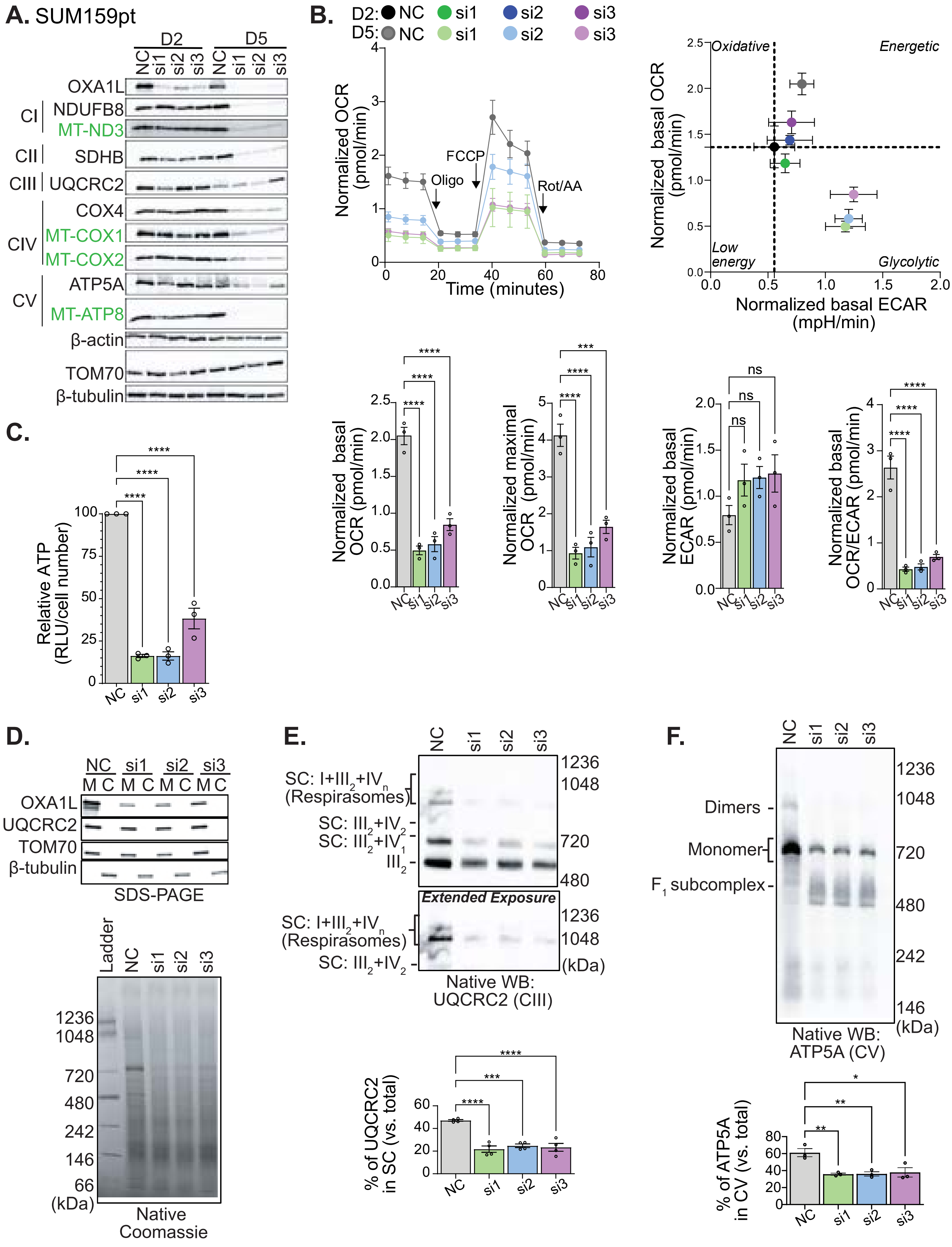

### Supplementary Figure 3

Supplementary Figure 3.

**A.**

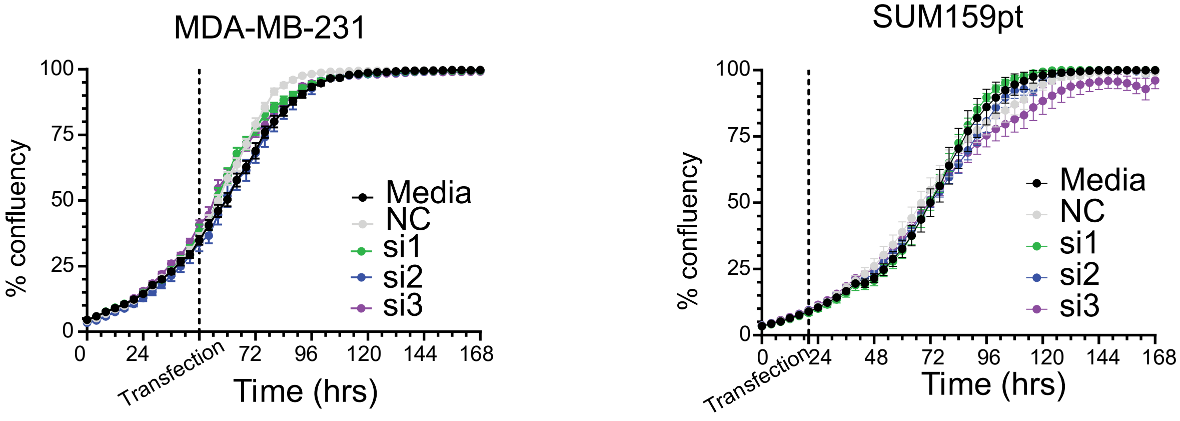


**B.**

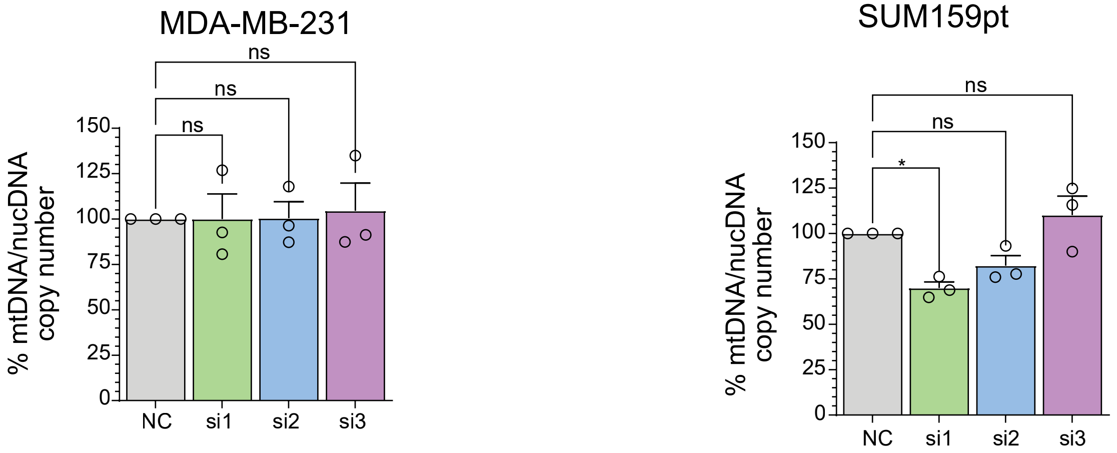


**C.**

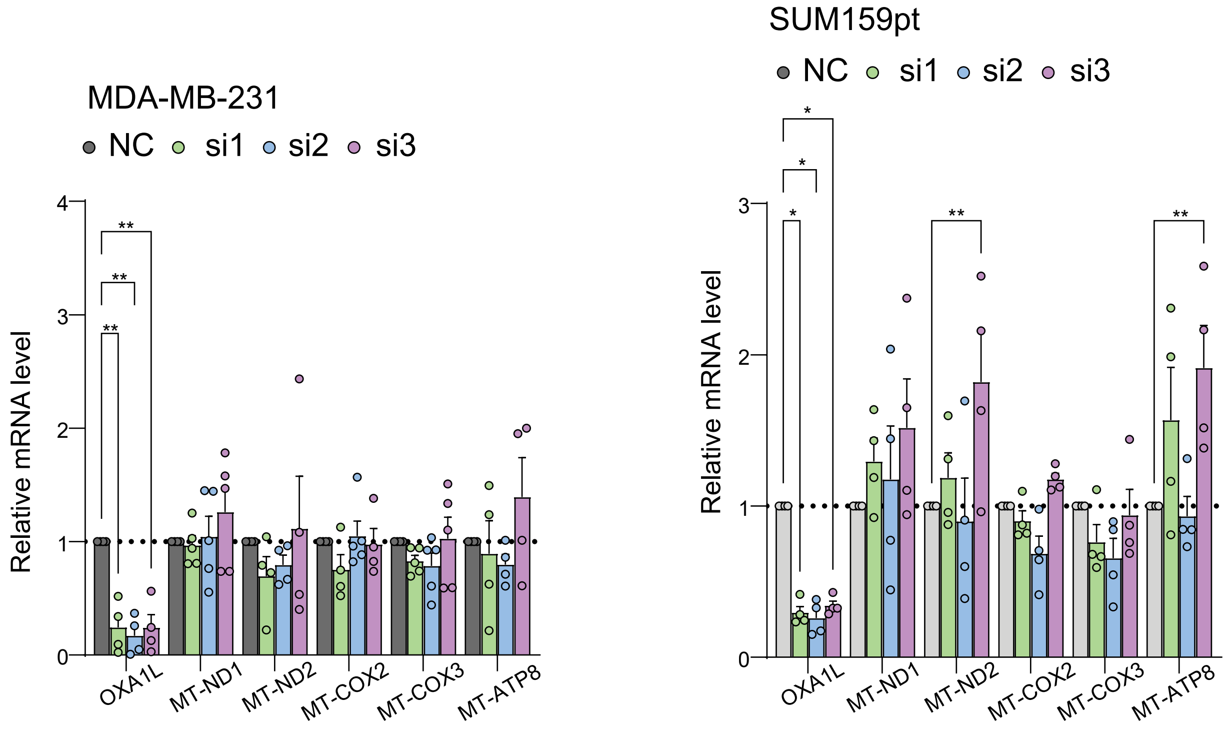

### Supplementary Figure 4

Supplementary Figure 4.

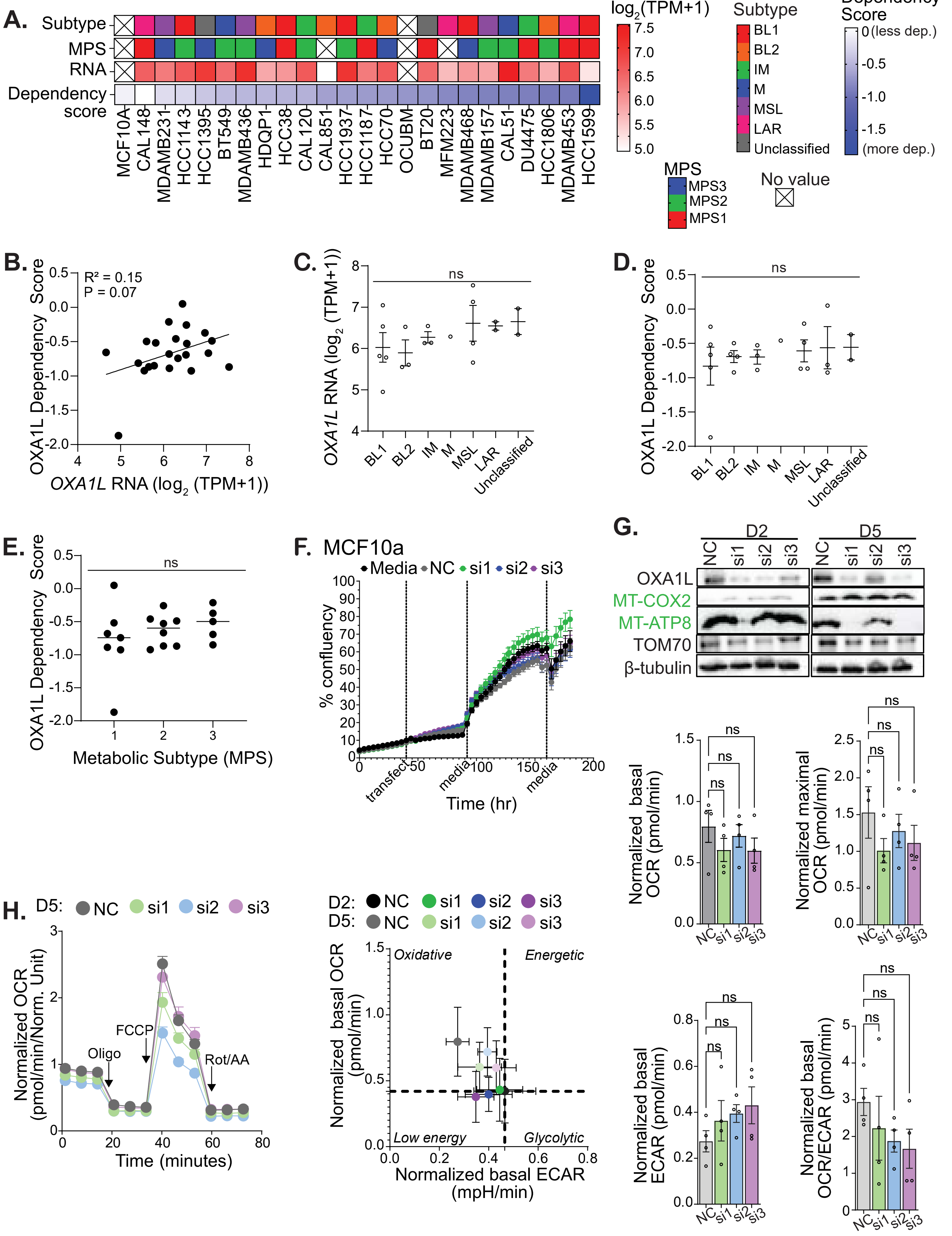

### Supplementary Figure 5

Supplementary Figure 5.

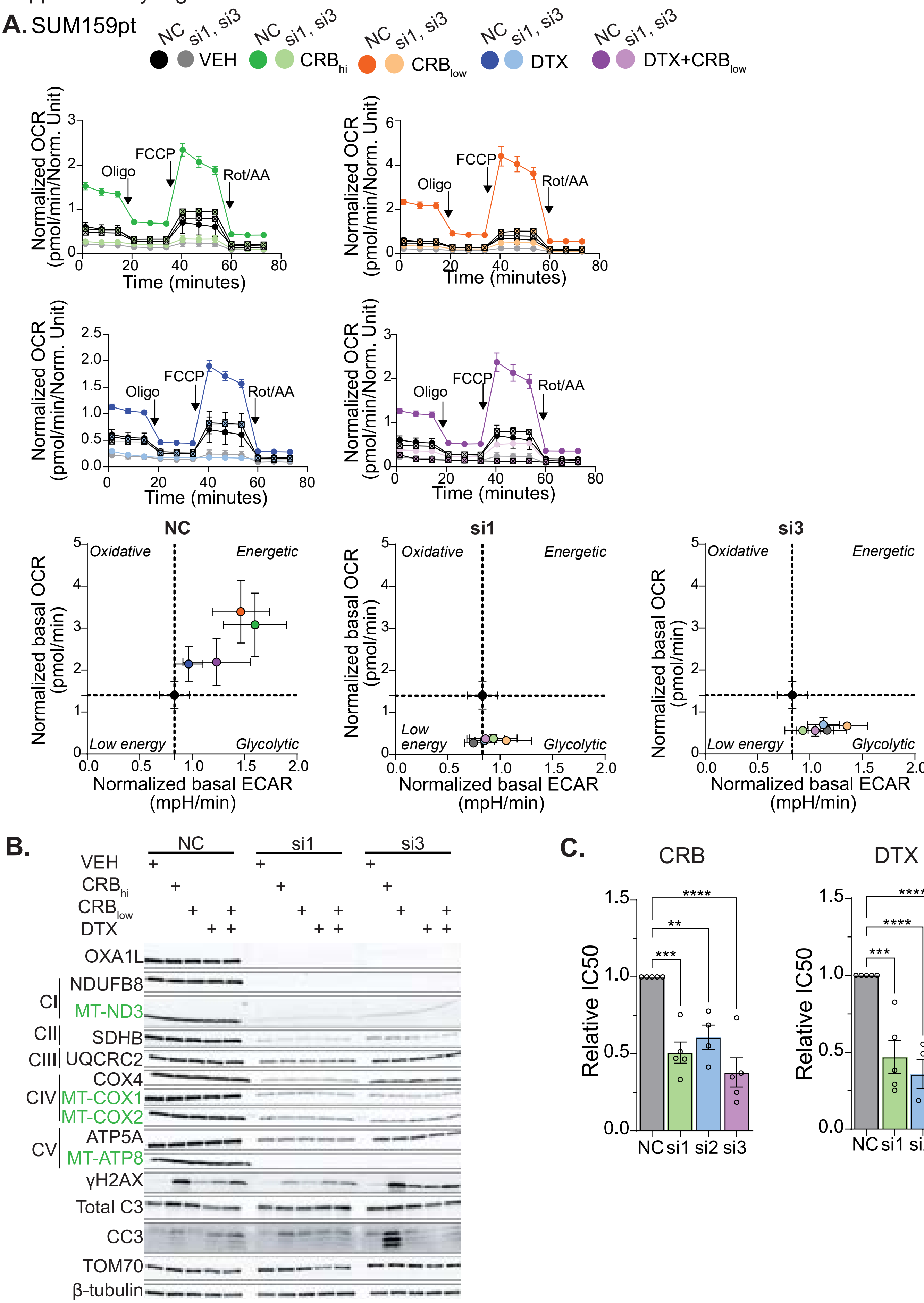

### Supplementary Figure 6

Supplementary Figure 6.

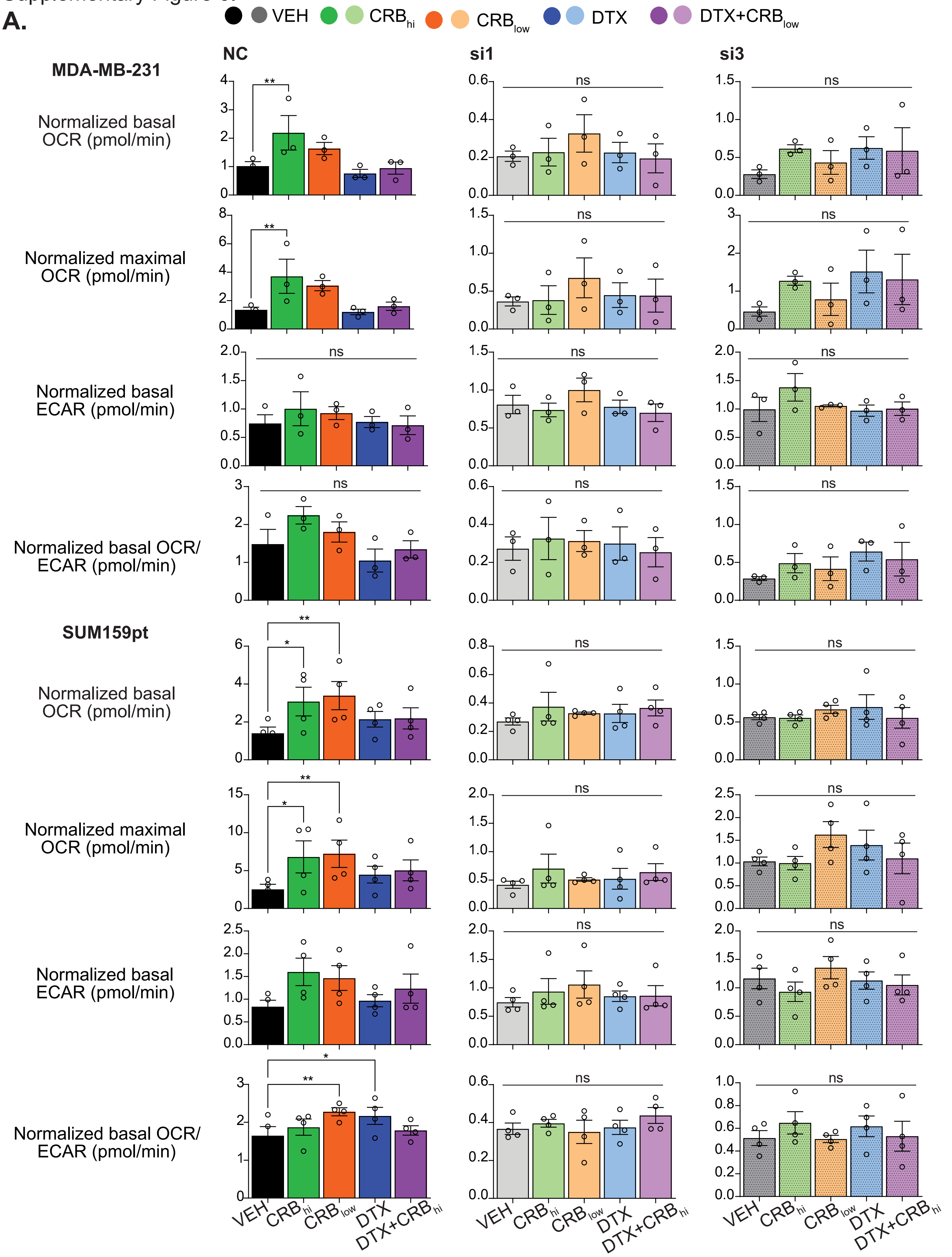

### Supplementary Figure 7

Supplementary Figure 7.

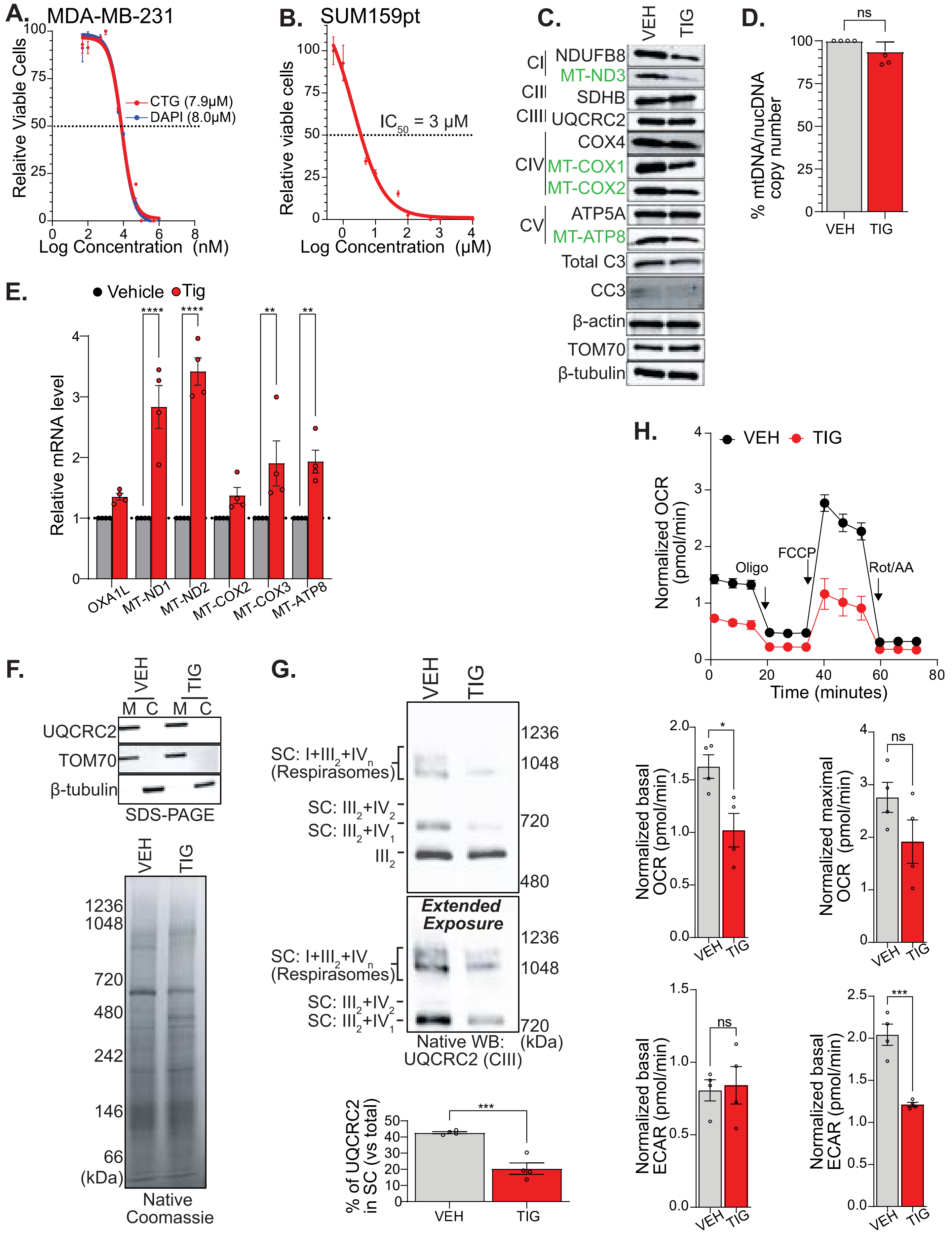

### Supplementary Figure 8

Supplementary Figure 8.

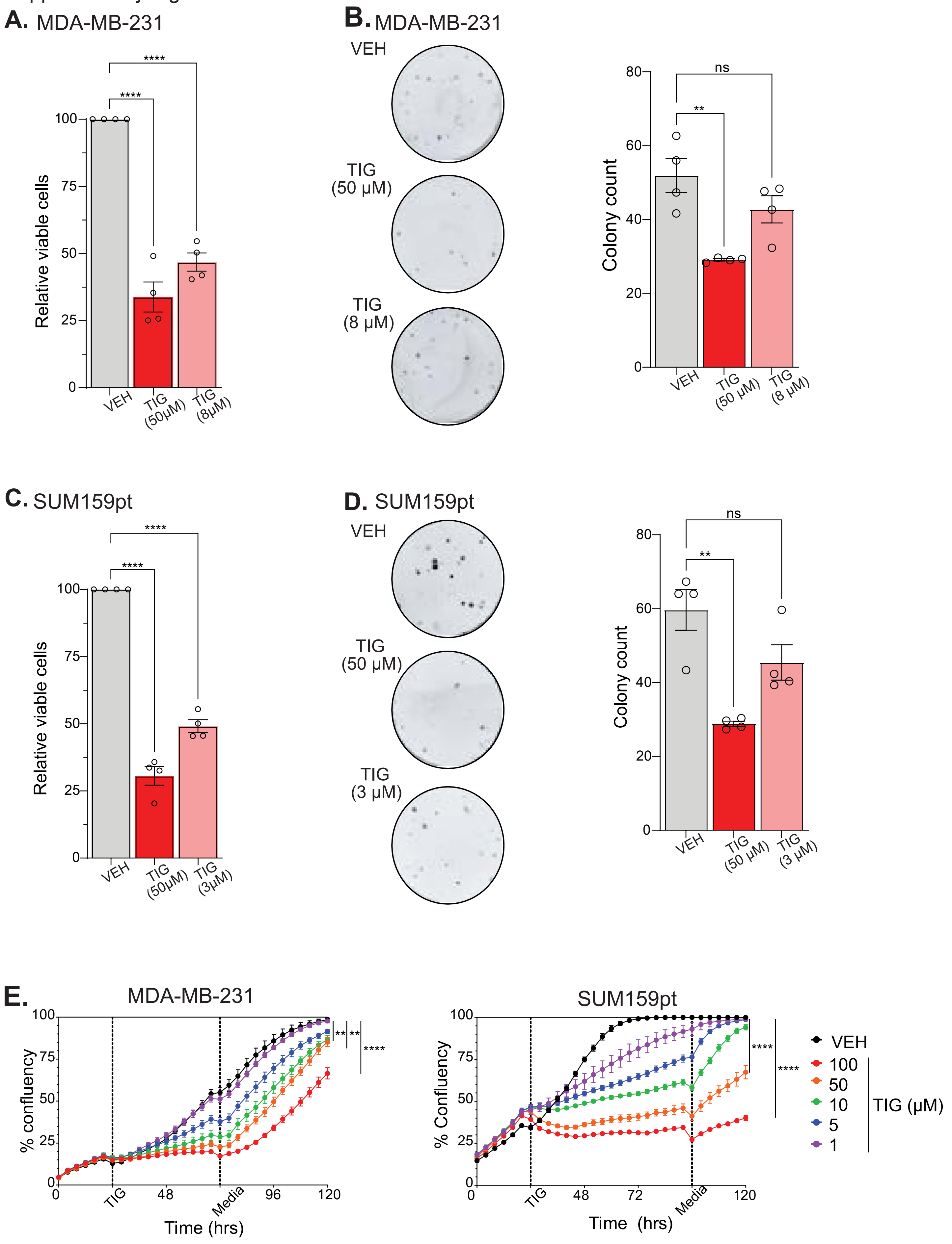

### Supplementary Figure 9

Supplementary Figure 9.

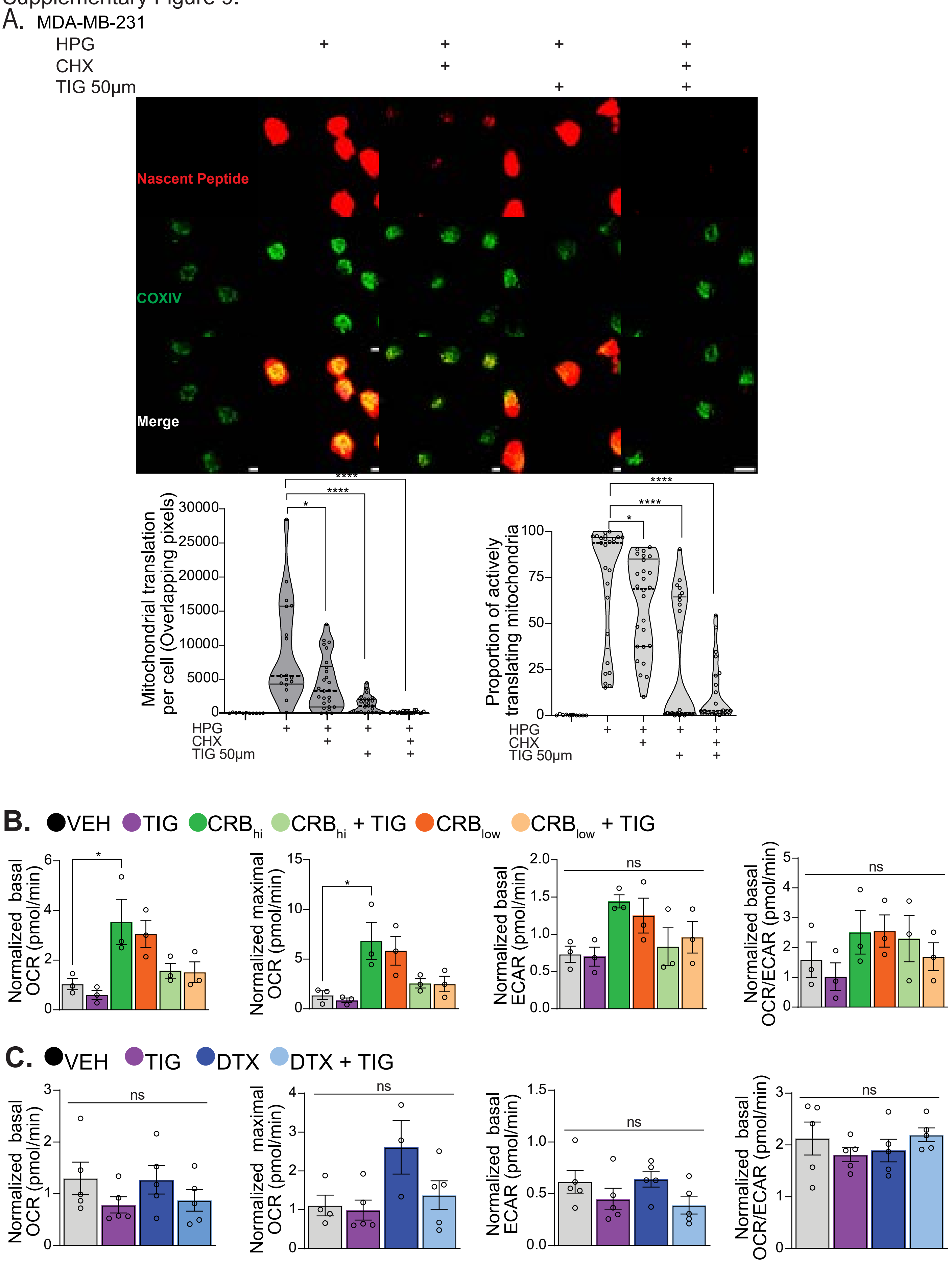

### Supplementary Figure 10

Supplementary Figure 10.

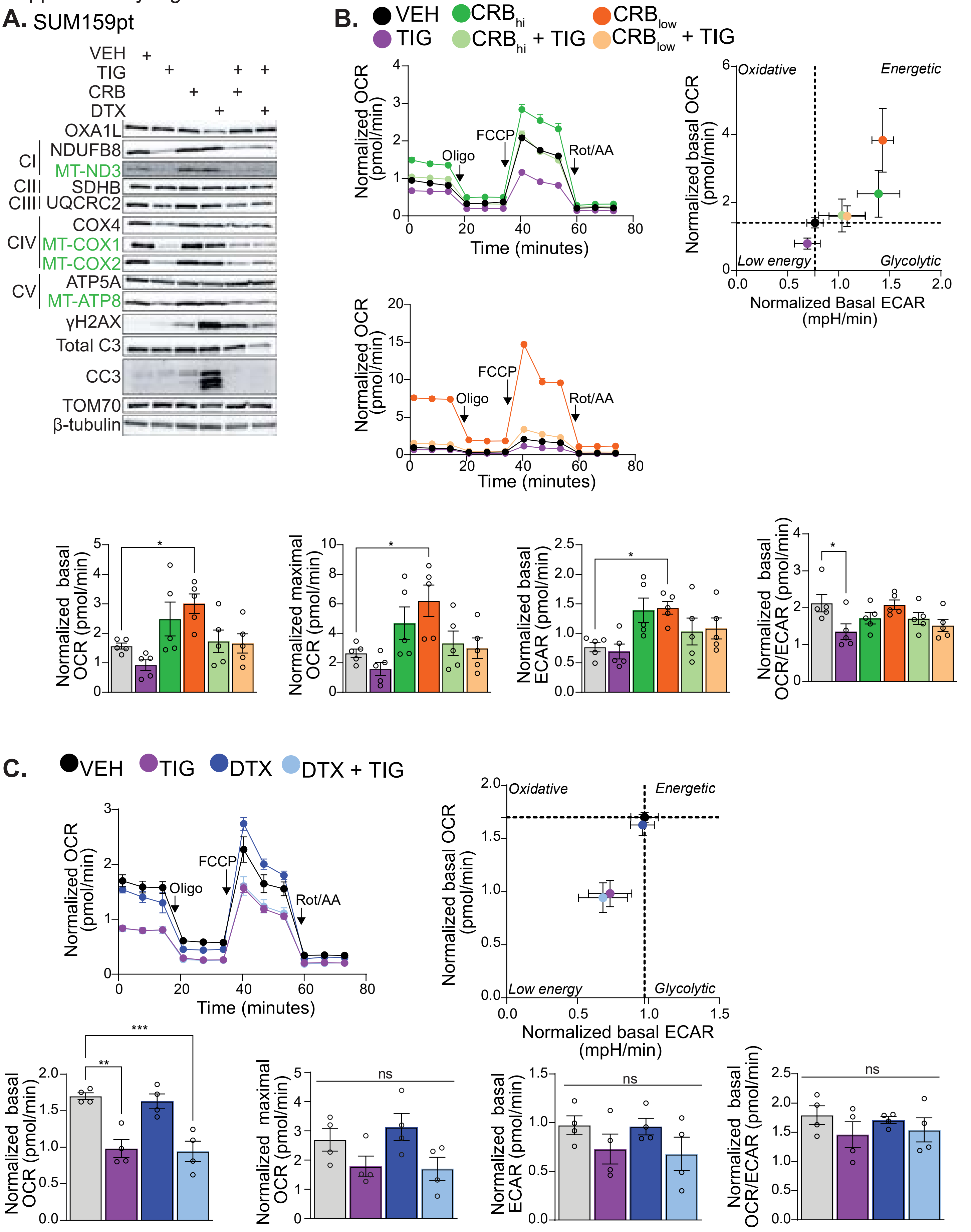

### Supplementary Figure 11

Supplementary Figure 11.

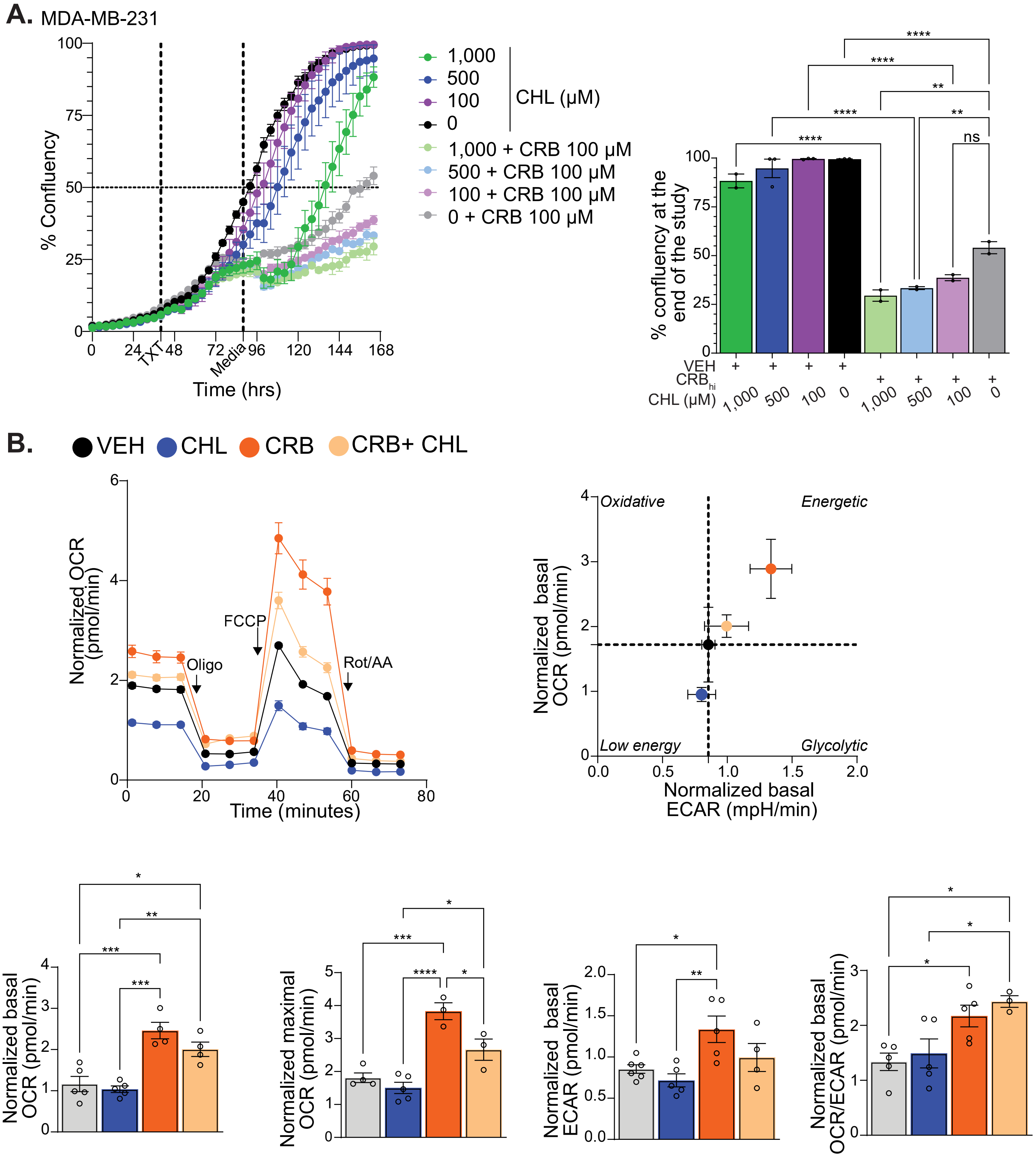

### Supplementary Figure 12

Supplementary Figure 12.

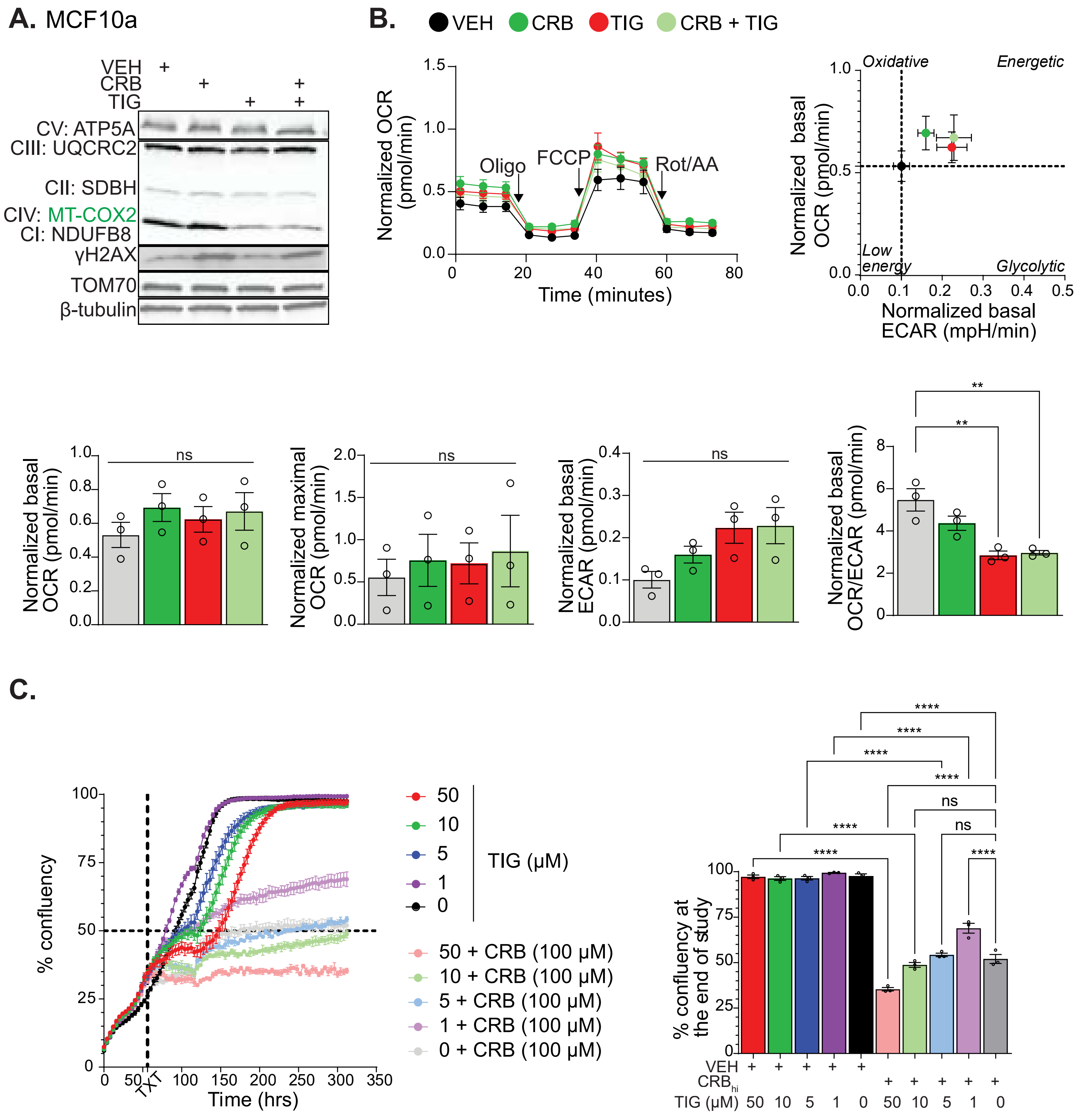
