## Supplementary Figure 13 for "Mitochondrial translation is a targetable dependency of chemo-refractory triple negative breast cancer"

A.

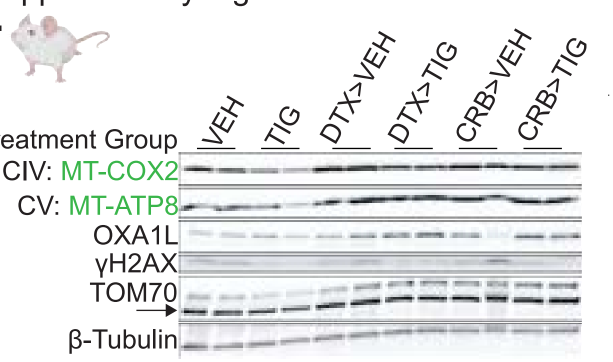


B.

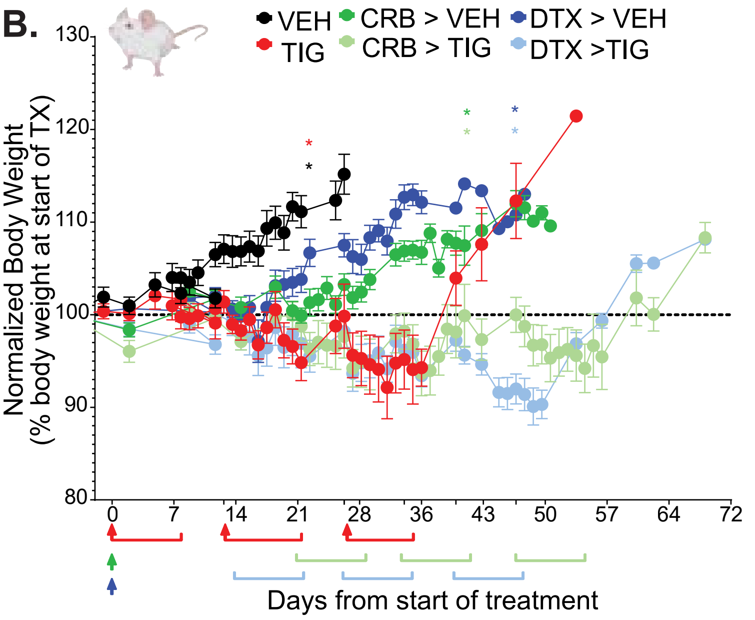


C.

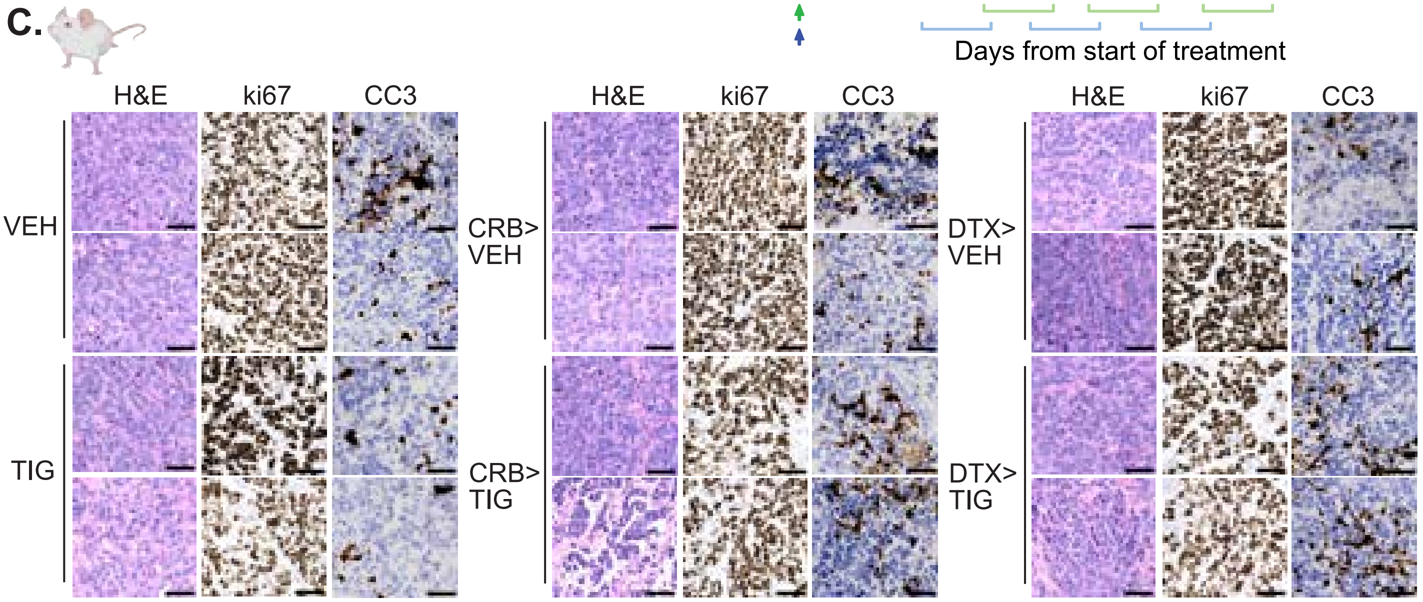


D.

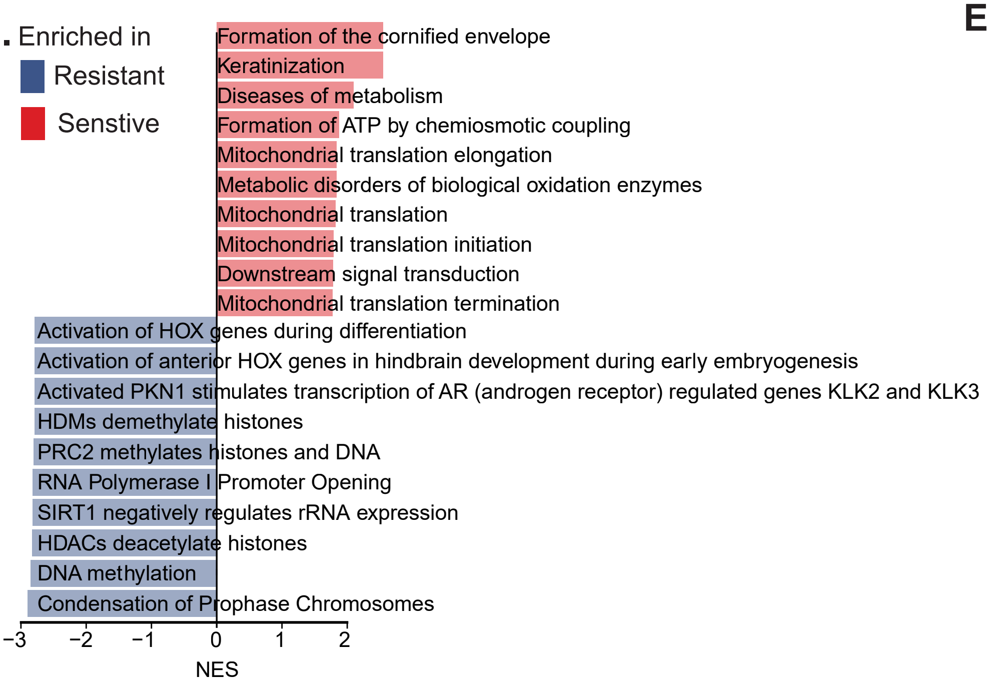


E.

| Rank | Leading edge proteins from mitochondrial translation pathway enriched in TIG sensitive spheroids |
|---|---|
| 1 | MRPS17 |
| 2 | GFM1 |
| 3 | MRPS18A |
| 4 | MRPL18 |
| 5 | MRPS11 |
| 6 | MRPL20 |
| 7 | MRPS25 |
| 8 | MRPS14 |
| 9 | MRPS2 |
| 10 | MRPS9 |
| 11 | MRPL48 |
| 12 | MRPS23 |
| 13 | TSFM |
| 14 | MRPL34 |
| 15 | MRPS28 |
| 16 | OXA1L |
| 17 | MRPL52 |
| 18 | MRPS21 |
| 19 | MRPS33 |
| 20 | MRPS15 |
